# Excitatory neurotransmission activates compartmentalized calcium transients in Müller glia without affecting lateral process motility

**DOI:** 10.1101/2021.09.02.458760

**Authors:** Joshua M. Tworig, Chandler Coate, Marla B. Feller

## Abstract

Neural activity has been implicated in the motility and outgrowth of glial cell processes throughout the central nervous system. Here we explore this phenomenon in Müller glia, which are specialized radial astroglia that are the predominant glial type of the vertebrate retina. Müller glia extend fine filopodia-like processes into retinal synaptic layers, in similar fashion to brain astrocytes and radial glia which exhibit perisynaptic processes. Using two-photon volumetric imaging, we found that during the second postnatal week, Müller glial processes were highly dynamic, with rapid extensions and retractions that were mediated by cytoskeletal rearrangements. During this same stage of development, retinal waves led to increases in cytosolic calcium within Müller glial lateral processes and stalks. These comprised distinct calcium compartments, distinguished by variable participation in waves, timing, and sensitivity to an M1 muscarinic acetylcholine receptor antagonist. However, we found that motility of lateral processes was unaffected by the presence of pharmacological agents that enhanced or blocked wave-associated calcium transients. Finally, we found that mice lacking normal cholinergic waves in the first postnatal week also exhibited normal Müller glial process morphology. Hence, outgrowth of Müller glial lateral processes into synaptic layers is determined by factors that are independent of neuronal activity.

## Introduction

Bidirectional signaling between neurons and glia is essential for circuit formation and function throughout the nervous system. Across developmental steps from neurogenesis to circuit maturation, glia monitor their environment and in turn regulate neuronal production, migration, and differentiation, promote synapse turnover, and regulate synaptic function via neurotransmitter uptake and ion buffering^1^. In the vertebrate retina, for example, Müller glia exhibit neurogenic potential^2–4^, promote circuit-specific wiring via secretion of synaptogenic molecules^5^, regulate phagocytosis of neuronal debris^6^, and limit neurotransmitter spillover via transporter activity^7^.

There is extensive evidence that neuronal signaling influences glial physiology in the adult brain. Individual astroglia extend fine processes that contact thousands of synapses and express an array of neurotransmitter receptors^8^. This enables glia to rapidly integrate neuronal activity, often involving intracellular calcium mobilization. Neuronal activity-evoked calcium events in glia range in size from small membrane-proximal microdomains to global cytosolic events mediated by various transmembrane proteins and calcium sources^9,10^. Pharmacological studies suggest compartmentalized and global calcium events are mediated by separate mechanisms and evoke different functional responses within glia, which lead to differential effects on neurotransmission^11–13^.

Here we use the mouse retina as a model to explore mechanisms and a possible function of neuron-glia signaling during development. In the retina, Müller glia are the predominant glial type, tiling the entire retinal space and interacting with every retinal cell type^14^. Similar to Bergmann glia of the cerebellum^15,16^, Müller glia exhibit a radial structure with a stalk extending from the soma, lateral processes extending from the stalk within synaptic layers, and endfeet contacting neuronal somata, axons, and vasculature. In the adult, this complex morphology enables Müller glia to mediate neurovascular coupling, maintain pan-retinal ion homeostasis, and modulate neuronal signaling in the adult ^17–19^. During both mouse^20,^ and zebrafish^21^ development, Müller glia undergo calcium transients in response to retinal waves, a term used to describe spontaneous bursts of depolarization that propagate across the retina. The functional relevance of wave-associated calcium signaling in Müller glia is not known.

One potential role for wave-associated calcium-signaling in Müller glia is in modulating outgrowth of lateral processes, which initiates and progresses during the same developmental window as retinal waves^14^. In other brain regions and model systems, neuronal activity-evoked calcium transients lead to morphological changes in glial processes. For example, in the hippocampus and somatosensory cortex, perisynaptic astrocytic processes undergo spatially localized, glutamate-evoked calcium transients which are essential for activity-evoked ensheathment of synapses^22^. In the cerebellum, Bergmann glial processes are highly motile during synaptogenic periods, and their ensheathment of Purkinje cells is impaired when glial glutamate transporters are knocked out^16,23^. Radial glia of the *Xenopus* optic tectum exhibit neuronal activity-evoked calcium transients and motility during visual system development^24^. In many of these systems, when glial motility is blocked, synaptic development, function, and plasticity are impaired^25^, highlighting the importance of glial dynamics in setting up and maintaining neural circuits.

Here, we combine morphological and calcium imaging, electrophysiology, and pharmacology to characterize Müller glial lateral process outgrowth during retinal waves and to determine the impact of neuronal activity on Müller glial morphology.

## Results

### Müller glial lateral process growth is non-uniform and dynamic across the IPL

Our goal is to determine whether spontaneous activity driven by retinal waves influences the morphological development of Müller glial cells. After their differentiation from retinal progenitor cells, which occurs during the first postnatal week, Müller glia extend processes laterally from their stalk into the inner plexiform layer (IPL)^26^. To visualize this process, we used the *GLAST-CreER;mTmG* reporter line to sparsely label Müller glia (Figure 1A). Lateral processes exhibited sublayer-specific outgrowth and distribution, and they reached an adult level of complexity soon after eye opening around postnatal day 14 (P14) (Figure 1B,D; see Figure 1-figure supplements 1 and 2 for statistical comparisons across sublayer and age, respectively). During the first days of Müller glia differentiation, lateral processes preferentially occupied the borders of the IPL (putative S1/S5), with most outgrowth occurring in the prospective ON half of the IPL (S3-S5) until about P10. By eye opening, after P12, processes arborized throughout the IPL and began to resemble their adult distribution, characterized by fewer processes in IPL sublayers S2/S4 than in S1/S3/S5. These observations are consistent with previous findings using the same mouse line and immunohistochemistry in fixed tissue^14^.

**Figure 1.**
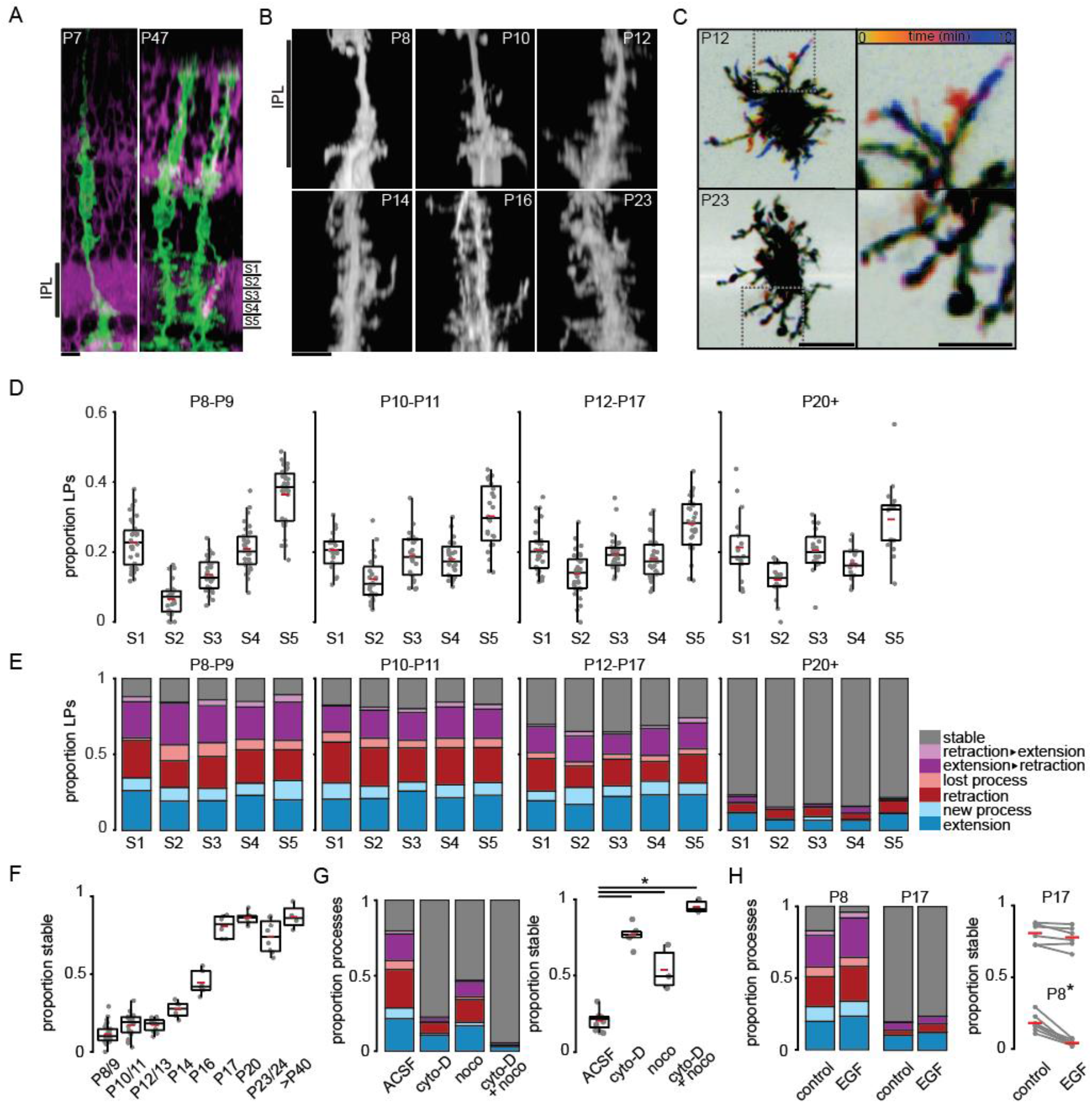
Müller glial lateral processes are highly motile during development. **(A)** Orthogonal projections of two-photon volumetric images of Müller glia expressing membrane-GFP (mGFP, green) in sparsely recombined *GLAST-CreER;mTmG* retinas at P7 (left) and P47 (right). Cells without Cre-mediated recombination express membrane-tdTomato (magenta). Scale bar 10 μm. IPL: inner plexiform layer. **(B)** Orthogonal projections of mGFP-expressing Müller glia from P8 to P23. Scale bar 10 μm. **(C)** Temporally color-coded projections of two-photon Z-stack time series showing motile processes at P12 (top) and stable processes at P23 (bottom). The location and dynamics of lateral processes at each time point can be determined by referencing the color time scale at bottom. Left scale bar 10 μm; right scale bar 5 μm. **(D)** Distribution of lateral processes across IPL sublayers through development. See Figure 1-figure supplement 1 for summary statistics and goodness-of-fit test statistics, and Figure 1-figure supplement 2 for tests for independent proportions across age groups. **(E)** Proportion of lateral processes in each sublayer that underwent extension (blue), sprouting (light blue), retraction (red), elimination (light red), a combination (purple), or that were stable (grey). **(F)** Proportion of total stable processes across development. See Figure 1-figure supplement 3 for statistical comparisons. **(G)** Proportion of total processes that were motile (left) and stable (right) in the absence and presence of blockers of cytoskeletal turnover. cyto-D: cytochalasin-D (5 μM); noco: nocodazole (10 μM). See Figure 1-figure supplement 4 for statistical comparisons. **(H)** Proportion of total processes that were motile (left) and stable (right) in the absence and presence of epidermal growth factor (EGF, 1 unit (100ng)/mL) at P8 and P17. See Figure 1-figure supplement 5 for statistical comparisons. * corresponds to p<0.05 for all comparisons. Source data available in Figure 1-source data 1.

During outgrowth from the primary stalks, we observed that Müller glial lateral processes across the IPL were highly motile. To characterize process motility, we conducted volumetric two-photon imaging of sparsely labeled processes at a rate of roughly 1 volume every 2 minutes for 10-minute epochs. We characterized motility that occurred during the imaging period by grouping events into distinct categories as follows: new processes that branched from stalks or from existing processes, extension of existing processes, retraction of processes without elimination, elimination of existing processes, or stable for processes that did not change length. We made several observations based on this analysis (Figure 1E). First, Müller glial cell lateral processes were highly motile across the IPL throughout the entire second postnatal week. Second, there was a slight bias toward new process sprouting rather than process loss. Third, we found there were roughly equal proportions of extending and retracting processes over 10-minute epochs of imaging, highlighting their rapid turnover during this period of development. Finally, we observed a sharp developmental transition toward stability a few days after eye opening (Figure 1C,E,F; Figure 1-figure supplement 3).

To assure that we could observe and quantify changes in process motility, we applied two manipulations. First, we slowed process motility with pharmacological agents that disrupt cytoskeletal proteins. Process motility was reduced by bath application of the microtubule-disrupting agent nocodazole (10 μM) and the actin polymerization inhibitor cytochalasin-D (5 μM), the combination of which led to processes stabilization (Figure 1G; Figure 1-figure supplement 4). Second, we enhanced process motility by bath application of epidermal growth factor (EGF; 1 unit (100ng)/mL; Figure 1H; Figure 1-figure supplement 5), which activates EGF receptors (EGFRs). EGFRs are expressed by Müller glia during the second postnatal week^27^ and increase motility in other cells via activation of Rac- and Rho-GTPase-dependent pathways^28,29^. Together, these results indicate that lateral process motility is facilitated by turnover of actin filaments and microtubules and may be modulated by growth factor signaling, pathways which potentially interact with neuronal activity^30,31^.

### Retinal waves activate compartmentalized calcium transients in Müller glia

Much of the morphological development of Müller glia occurs when retinal waves are present (Figure 1 and [14]). Furthermore, previous studies have demonstrated that retinal waves induce increases in intracellular calcium in Müller glia^20,21^. However, these studies did not assess whether there are distinct calcium compartments within Müller glia which could potentially have distinct impacts on lateral process motility. The two compartments in Müller glia we focused on were the central stalks and the lateral processes within the IPL.

We conducted simultaneous two-photon calcium imaging of Müller glial stalks and processes and voltage clamp recordings from retinal ganglion cells (RGCs). Several strategies for calcium imaging from Müller glia were used. First, retinas were isolated from *GLAST-CreER; cyto-GCaMP6f* mice, which express genetically encoded calcium indicator selectively in Müller glia. Second, a subset of retinas were isolated from WT mice and bath-loaded with the chemical calcium dye Cal520, which also selectively labels Müller glia^20,32^. Using these two approaches, we were able to clearly identify the boundaries of Müller glial stalks, while calcium in lateral processes was assessed using regions of interest (ROIs) within the areas of IPL intervening glial stalks (Figure 2A). Third, retinas were isolated from mice expressing membrane-bound calcium reporter in glial cells (Lck-GCaMP6f in *GLAST-CreER* mice)^33^, which enabled resolution of lateral processes but poor detection of cytosolic calcium transients within stalks (Figure 2-figure supplement 1). Using all of these approaches, we verified that Müller glial stalks and lateral processes exhibited calcium transients in response to retinal waves as previously reported^20^, as well as spontaneous, non-wave-associated calcium transients.

**Figure 2.**
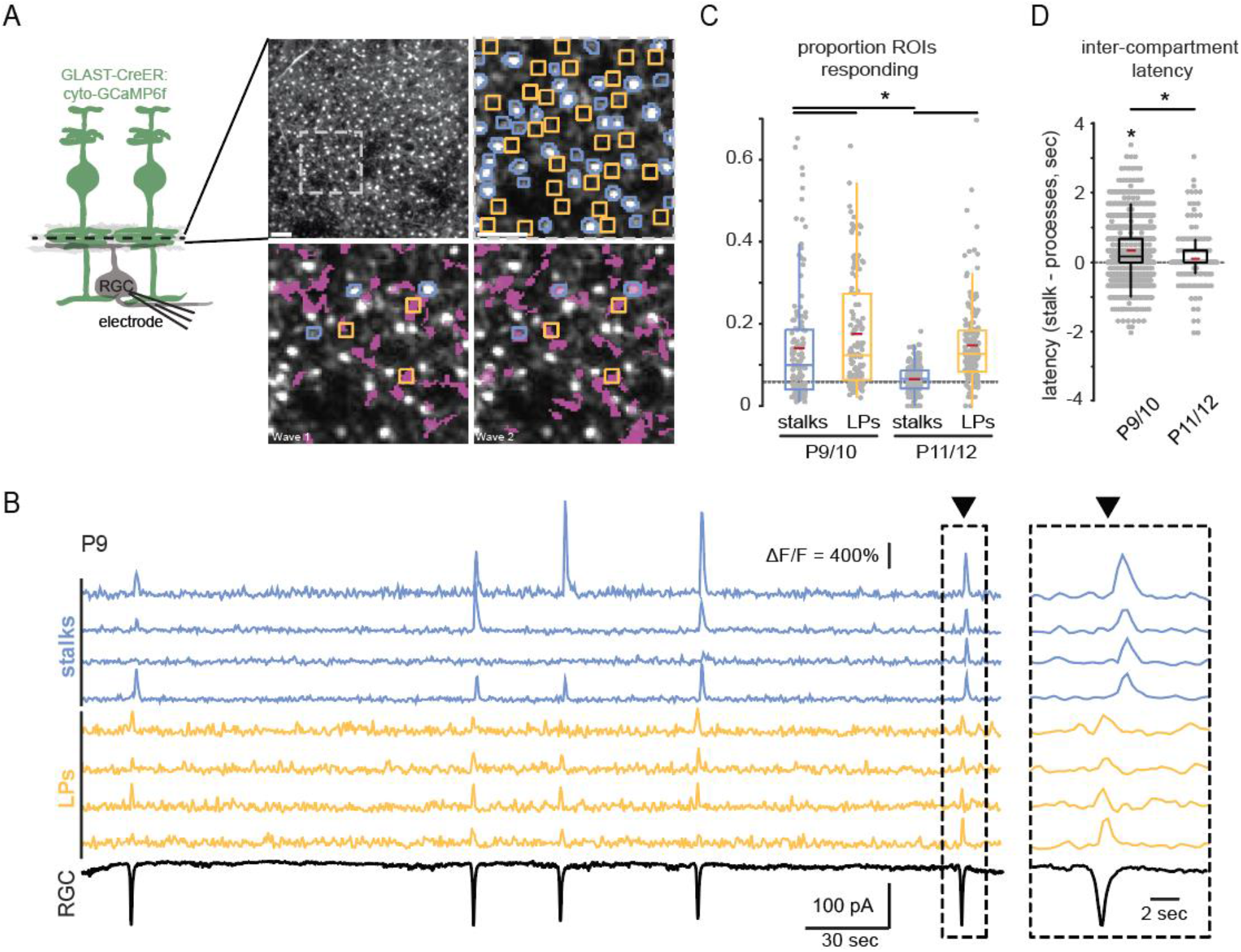
Retinal wave-associated calcium transients are compartmentalized within glial stalks and lateral processes. **(A)** *Left,* diagram of experimental setup; cytosolic GCaMP6f is expressed in Müller glia (green) via GLAST-CreER, and a two-photon microscope is used to image calcium transients in glial stalks and lateral processes in the IPL (dashed line indicates focal depth in whole mount retina). Excitatory currents are recorded from a retinal ganglion cell (RGC) to detect retinal waves. *Right*, average projection of a field of view (FOV) showing GCaMP6f-expressing Müller glia in retinal whole mount. Upper left panel is the full FOV; upper right panel is a magnified portion of this FOV overlayed with regions of interest (ROIs) for stalks (blue) and lateral processes (yellow); bottom panels show the extent of glial activation by two retinal waves (active regions overlayed in magenta). **(B)** Example traces of fractional change in fluorescence of GCaMP6f-expressing Müller glia showing wave-associated calcium transients in stalks (blue) and processes (yellow) and wave-associated EPSCs recorded from an RGC (black). *Right,* glial calcium response during a retinal wave, expanded in time to highlight temporal delay of stalk transients relative to lateral process transients. Black trace is a voltage-clamp recording from an RGC held at −60mV. **(C)** Proportion of total stalk and lateral process ROIs that respond to retinal waves at P9/10 and P11/12. Grey dashed line indicates the proportion of ROIs undergoing spontaneous calcium transients at randomly selected times. See Figure 2-figure supplement 2 for summary statistics. **(D)** Intercompartment response latency, calculated as latency of stalk responses from the median lateral process response time for retinal waves at P9/10 and P11/12. See Figure 2-figure supplement 3 for summary statistics. Image scale bars 10μm. Source data available in Figure 2-source data 1.

We observed several differences in wave-associated calcium transients between stalks and lateral processes. Wave-associated stalk transients propagated throughout the stalk and involved many processes, which we refer to as a global transient. Lateral processes participated in some global transients but also exhibited wave-associated calcium transients independent of stalks (Figure 2A,B and Figure 2-figure supplement 1). Hence, lateral processes and stalks differentially participated in waves, with a greater proportion of lateral process ROIs responding to waves than stalk ROIs. In addition, wave-associated calcium transients in lateral processes were consistently observed from P9 to P12, while those in stalks were significantly reduced in terms of ΔF/F amplitude and proportion of stalks participating by P11/12 (Figures 2C, 3A; Figure 2-figure supplement 2). Finally, the subset of stalks that responded to retinal waves did so with a slight delay relative to lateral processes (Figure 2B,D; Figure 2-figure supplement 3). Hence, Müller glial stalks and lateral processes can undergo spatiotemporally distinct calcium signaling events.

We next sought to test whether wave-associated calcium transients in stalks vs. lateral processes are mediated via activation of distinct signaling pathways. Muscarinic acetylcholine receptors (mAChRs) have been implicated in Müller glial response to retinal waves,^20^ and isolated Müller glia undergo M1 mAChR-dependent responses to cholinergic agonists^34^. Bath application of pirenzepine (5 μM), a selective antagonist of M1 mAChRs, led to a significant reduction in wave-associated calcium transients in stalks, while wave-associated transients remained in lateral processes (Figure 3A). This effect was age-dependent, as wave-associated transients in glial stalks had greater sensitivity to M1 mAChR block at P9/10 than at P11/12 (Figure 3B,C; Figure 3-figure supplements 1–4). Interestingly, we occasionally detected wave-associated stalk calcium transients that were insensitive to pirenzepine. Pirenzepine-insensitive stalk responses exhibited similar amplitude and latency to those of lateral processes (Figure 3C,D; Figure 3-figure supplement 5). These data suggest that wave-associated responses in stalks are controlled by multiple mechanisms in addition to activation of M1 mAChRs. Note, pirenzepine had minimal impact on the amplitude and frequency of compound excitatory postsynaptic currents (EPSCs) and inter-wave-interval (IWI, Figure 3E; Figure 3-figure supplement 6), indicating pirenzepine had a selective effect on Müller glial M1 mAChRs, rather than on wave-generating circuits.

**Figure 3.**
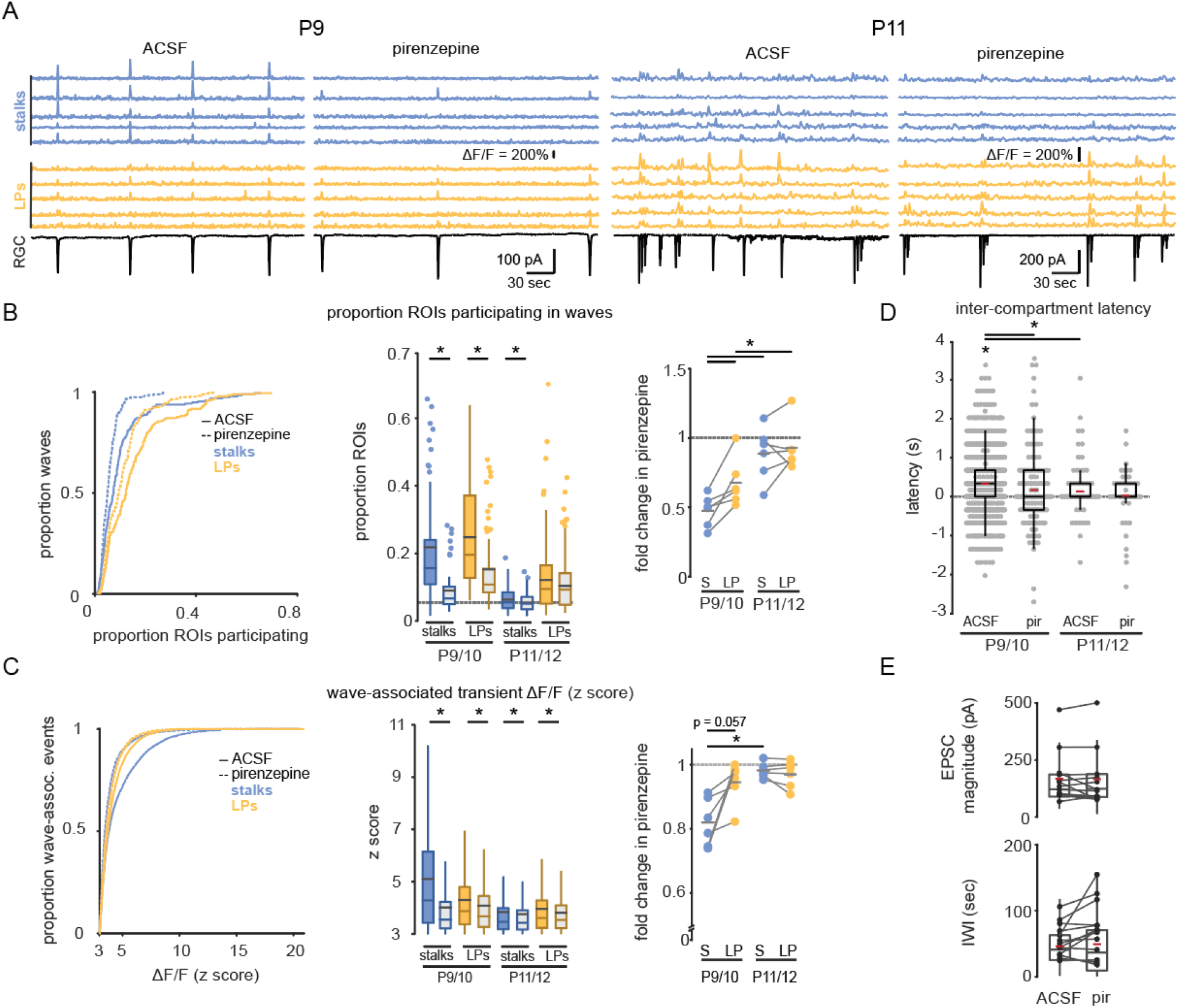
M1 mAChRs mediate retinal wave-associated calcium transients in glial stalks but not in lateral processes. **(A)** Example traces of fractional change in fluorescence of GCaMP6f-expressing Müller glia (top) showing wave-associated calcium transients in stalks (blue) and processes (yellow), and wave-associated EPSCs recorded from a retinal ganglion cell (RGG, bottom) in the absence and presence of pirenzepine (5 μM) at P9 (left) and P11 (right). **(B)** *Left*, cumulative distribution of proportion of stalks and lateral processes that exhibited wave-associated calcium transients in absence (solid line) and presence of pirenzepine (dotted line). *Middle*, box plots showing proportion of ROIs that exhibited wave-associated calcium transients. Grey dashed line indicates the proportion of ROIs that exhibited spontaneous calcium transients at randomly selected times. See Figure 1-figure supplement 1 for summary statistics. *Right,* fold-change in proportion of ROIs that exhibited wave-associated calcium transients in presence of pirenzepine. See Figure 1-figure supplement 2 for summary statistics. **(C)** *Left*, cumulative distribution of the ΔF/F amplitude of wave-associated calcium transients in the absence and presence of pirenzepine. See Figure 1-figure supplement 3 for summary statistics. *Middle*, box plots showing ΔF/F amplitudes separated by age. *Right,* fold-change in wave-associated ΔF/F amplitude in pirenzepine. See Figure 1-figure supplement 4 for summary statistics. **(D)** Intercompartment response latency in each age and condition. See Figure 1-figure supplement 5 for summary statistics. **(E)** Summary data for magnitude (top) and interval between (bottom) compound excitatory post-synaptic current (EPSC) amplitude associated with retinal waves. See Figure 1-figure supplement 6 for summary statistics. Source data available in Figure 3-source data 1.

To confirm that M1 mAChR signaling preferentially impacts wave-associated stalk transients, we enhanced acetylcholine (ACh) release during waves by bath-applying the GABAA receptor antagonist gabazine (5 μM)^35^. As expected, the presence of gabazine increased the amplitude of EPSCs recorded from RGCs during retinal waves (Figure 4E; Figure 4-figure supplement 6). Gabazine also increased the proportion of stalks and lateral process ROIs that underwent wave-associated calcium transients (Figure 4A,B; Figure 4-figure supplement 1), but with differential compartment-specific effects: there was a significantly larger gabazine-induced fold change in proportion of stalks compared with lateral processes responding to waves, and gabazine led to a nearly 2-fold increase in the amplitude of wave-evoked fluorescence change in stalks, while only slightly increasing the response amplitude in lateral processes (Figure 4C; Figure 4-figure supplements 2–4). This effect was independent of age. Subsequent application of pirenzepine reduced the extent, magnitude, and latency of stalk responses to waves while largely sparing lateral process responses (Figure 4B-D; Figure 4-figure supplements 1–5) and without altering wave properties (Figure 4E; figure supplement 6). Taken together, these data indicate that M1 mAChRs mediate wave-associated global calcium transients in Müller glial stalks and support the conclusion that stalks and lateral processes are functioning as distinct calcium compartments.

**Figure 4.**
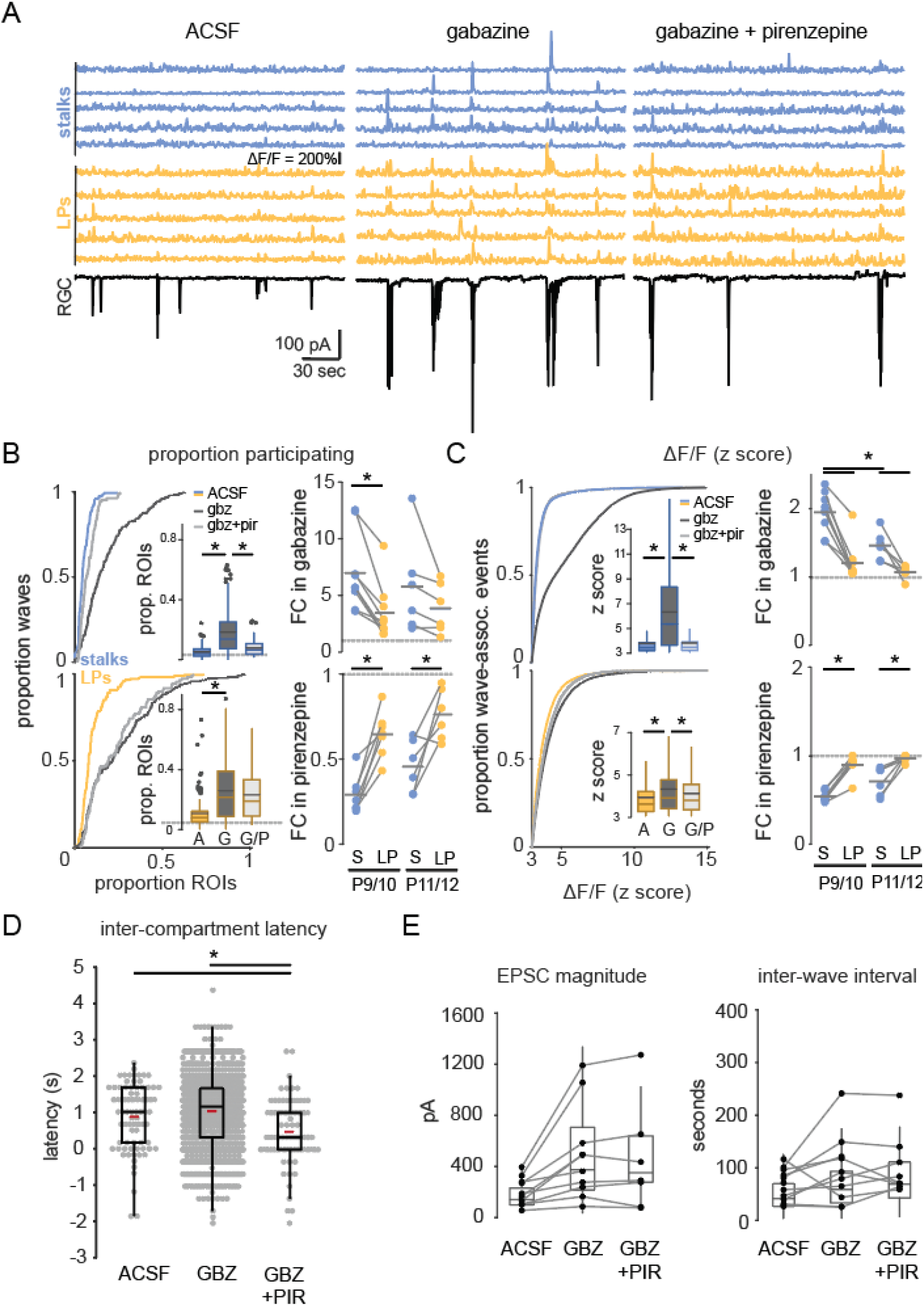
GABAA receptor block potentiates wave responses in stalks via M1 mAChR activation. **(A)** Examples traces of fractional change in fluorescence of GCaMP6f-expressing Müller glia showing wave-associated calcium responses (top) and wave-associated EPSCs recorded from an RGC (bottom) in control (left), 5 μM gabazine (middle), and gabazine plus 5 μM pirenzepine (right). **(B)** *Left*, cumulative distributions and inset box plots of proportion of stalk ROIs (top) and lateral process ROIs (bottom) that exhibited wave-associated calcium transients in control (solid line, A), gabazine (dotted line, G), and gabazine/pirenzepine (dashed line, G/P). Grey dashed line in box plots indicates the proportion of ROIs undergoing spontaneous calcium transients at randomly selected times. *Right,* fold-change (FC) in proportion ROIs responding in gabazine vs. ACSF (top) and in pirenzepine vs. gabazine (bottom). Grey dashed line indicates fold-change of 1. See Figure 4-figure supplements 1 and 2 for summary statistics. **(C)** *Left,* cumulative distributions of normalized wave-associated calcium transient amplitudes in ACSF, gabazine, and gabazine/pirenzepine, with inset box plots generated from the same data. *Right,* fold-change in normalized calcium response amplitude after gabazine (top) and pirenzepine (bottom). See figure 4-figure supplements 3 and 4 for summary statistics. **(D)** Intercompartment latency in wave-associated calcium transients in control, gabazine (gbz), and gabazine/pirenzepine (gbz+pir). See figure 4-figure supplement 5 for summary statistics. **(E)** Summary data for magnitude (top) and interval between (bottom) compound excitatory post-synaptic current (EPSC) associated with retinal waves in control, gabazine, and gabazine/pirenzepine for paired FOVs. See figure 4-figure supplement 6 for summary statistics. Source data available in Figure 4-source data 1.

### Müller glial process motility occurs independent of wave-associated calcium transients

Is Müller glial lateral process motility impacted by neuronal activity? To test whether neuronal activity influences Müller glial lateral process motility, we assessed the impact of multiple pharmacological agents on process motility after the first postnatal week (Figure 5). First, we assessed the impact of agents that modulated mAChR signaling resulting from neural activity (Figure 5A). Gabazine, which potentiated calcium transients in glial processes and stalks, in part via activation of M1 mAChRs, did not change the proportion of processes undergoing motility. Pirenzepine, which blocks M1 mAChRs and specifically abolished wave-associated calcium transients in glial stalks, led to a slight increase in proportion of processes exhibiting stability, but this did not reach statistical significance in paired tests (Figure 5A,F; see Figure 5-figure supplement 1 for summary statistics). Second, we assessed the impact of glutamatergic signaling, which is also implicated in wave-associated Müller glial calcium transients^20,21^. TBOA, which blocks glutamate transporters and enhances wave-associated calcium transients in Müller glia by increasing glutamate spillover^20,36^, also led to a small but non-statistically significant increase in the proportion of stable processes (Figure 5B,F). In contrast, blocking retinal waves with a combination of DNQX and AP5, which also reduced calcium transients in Müller glia (Figure 3-figure supplement 7), had no impact on motility (Figure 5B,F). These observations together indicate that lateral process motility does not arise from wave-associated calcium transients in stalks or processes.

**Figure 5.**
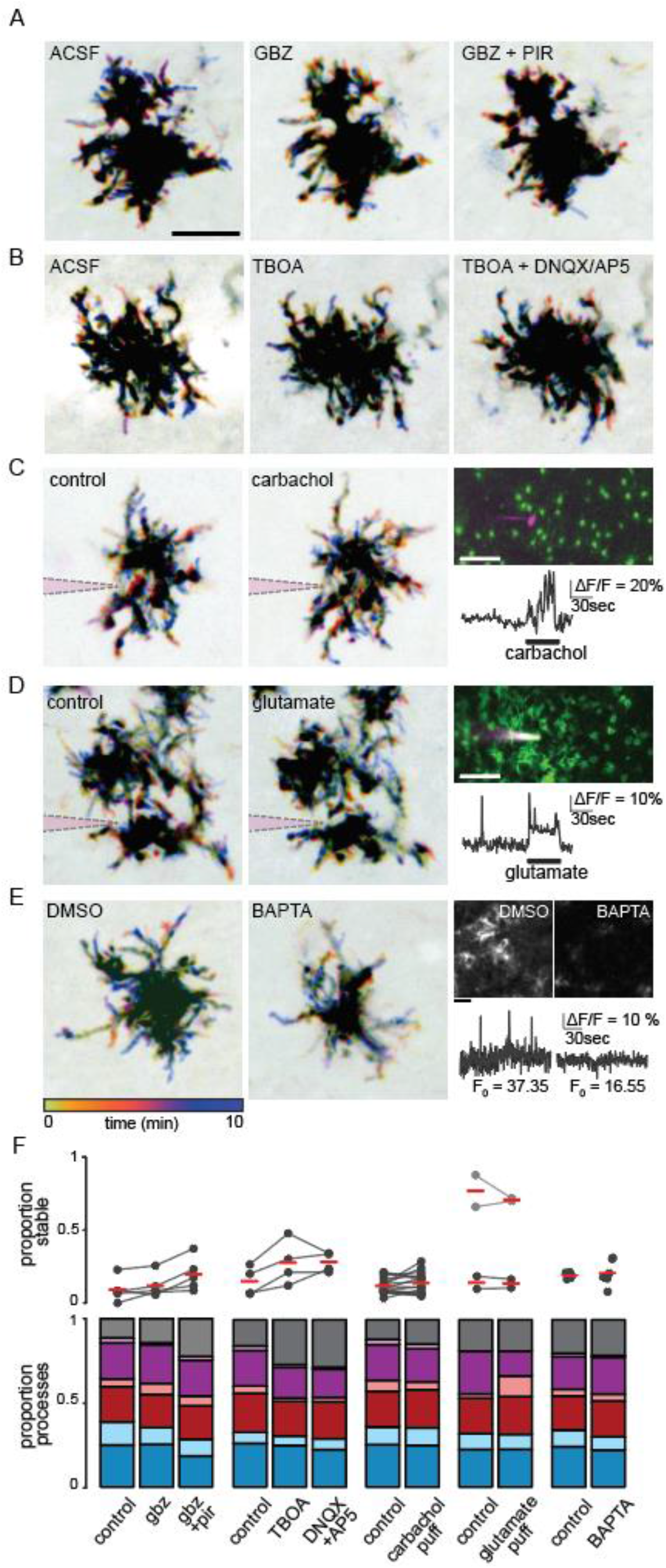
Müller glial lateral process motility is unaffected by manipulations of retinal waves. **(A)** Temporally color-coded projections of two-photon Z stack time series showing motile and stable processes in control (ACSF), gabazine (5 μM), and gabazine plus pirenzepine (5 μM). Scale bar 10 μm. **(B)**Same as A in control (ACSF), TBOA (25 μM), and TBOA plus DNQX (20 μM) and AP5 (50 μM) as pharmacological manipulations of retinal waves. **(C)** and **(D)** show motility (left, middle) and calcium responses (right, scale bar 20 μm) to iontophoretically applied carbachol and glutamate. A sharp electrode (indicated in magenta) was filled with agonist and current continuously applied to eject agonist into the IPL near mGFP-expressing processes to image motility, or near GCaMP6f-expressing processes to image calcium responses. **(E)** *Left,* motility persisted even when intracellular calcium was chelated via BAPTA-AM bath application, which significantly reduces baseline calcium and eliminates neuronal activity. Scale bars 10μm. *Right,* average projections (top) and normalized traces of GCaMP6f fluorescence (bottom) in DMSO and BAPTA. **(F)** Summary data showing proportion of total processes exhibiting stability (top) and other categories of motility (bottom) for drug conditions shown in A-E. See Figure 5-figure supplement 1 for summary statistics. Source data available in Figure 5-source data 1.

To further explore a role for ACh and glutamate in influencing motility, we tested whether local release of neurotransmitter would alter the motility of nearby processes, as is the case for perisynaptic astrocytic processes elsewhere in the brain, where it is postulated that local elevations in neurotransmitter promote process growth and synapse coverage^22,37^. We locally perfused via continuous iontophoresis the AChR agonist carbachol or glutamate onto glial processes in *GLAST-CreER;mTmG* retinas with sparse Cre-mediated recombination. In parallel experiments using retinas from *GLAST-CreER;GCaMP6f* mice, we confirmed that this method induced strong calcium transients in stalks and lateral processes. Despite this, we observed no impact of local perfusion of agonists on glial cell lateral process motility (Figure 5C,D,F).

As a final test for whether intracellular calcium signaling in Müller glia is required for motility, we assessed motility after bath-loading the potent calcium chelator, BAPTA-AM. Despite blocking retinal waves and all calcium transients in Müller glia, the presence of BAPTA did not alter the proportion of processes exhibiting motility. Together these data suggest that motility of Müller glial lateral processes persists in absence of intracellular calcium transients or neuronal activity (Fig. 5E,F).

### Long-term blockade of cholinergic retinal waves does not alter Müller glial morphology

Thus far, we have assessed the role of neuronal activity and calcium signaling on Müller glial lateral process motility using live imaging during the second postnatal week, while there is lateral process outgrowth. However, during the first postnatal week, there are wave-associated calcium transients in Müller glia also mediated by activation of mAChRs^20^. To test whether retinal wave-associated calcium signaling during this first postnatal week influences later process outgrowth during the second postnatal week, we assessed glial morphology in retinas isolated from mice lacking the β2 nicotinic acetylcholine receptor subunit (β2-nAChR-KO). β2-nAChR-KO mice exhibit significantly reduced cholinergic retinal waves^38–40^ and therefore have reduced M1 mAChR-induced signaling in glial cells.

Morphology of individual Müller glial cells was assessed by filling glia with Alexa-488 via sharp pipette in P12 and P30 wild type and β2-nAChR-KO mice. Filled cells were visualized via 2-photon volumetric imaging in live tissue, and processes were traced for subsequent morphological analysis (Figure 6A-C). At both P12 and P30, individual Müller cells exhibited lateral processes throughout the IPL, with more processes in the ON-half of the IPL than in the OFF-half prior to eye opening, in agreement with Figure 1. Detailed assessment shows that by P12, Müller glial lateral processes exhibited nearly the same level of complexity as observed in the adult. In addition, we found no significant difference between wild type control and β2-nAChR-KO Müller glia in terms of several morphological parameters including complexity, number, area, and length of lateral processes (Figure 6D,E; Figure 6-figure supplement 1). These data suggest that cholinergic signaling from retinal waves does not play a direct role in morphological development of glial processes.

**Figure 6.**
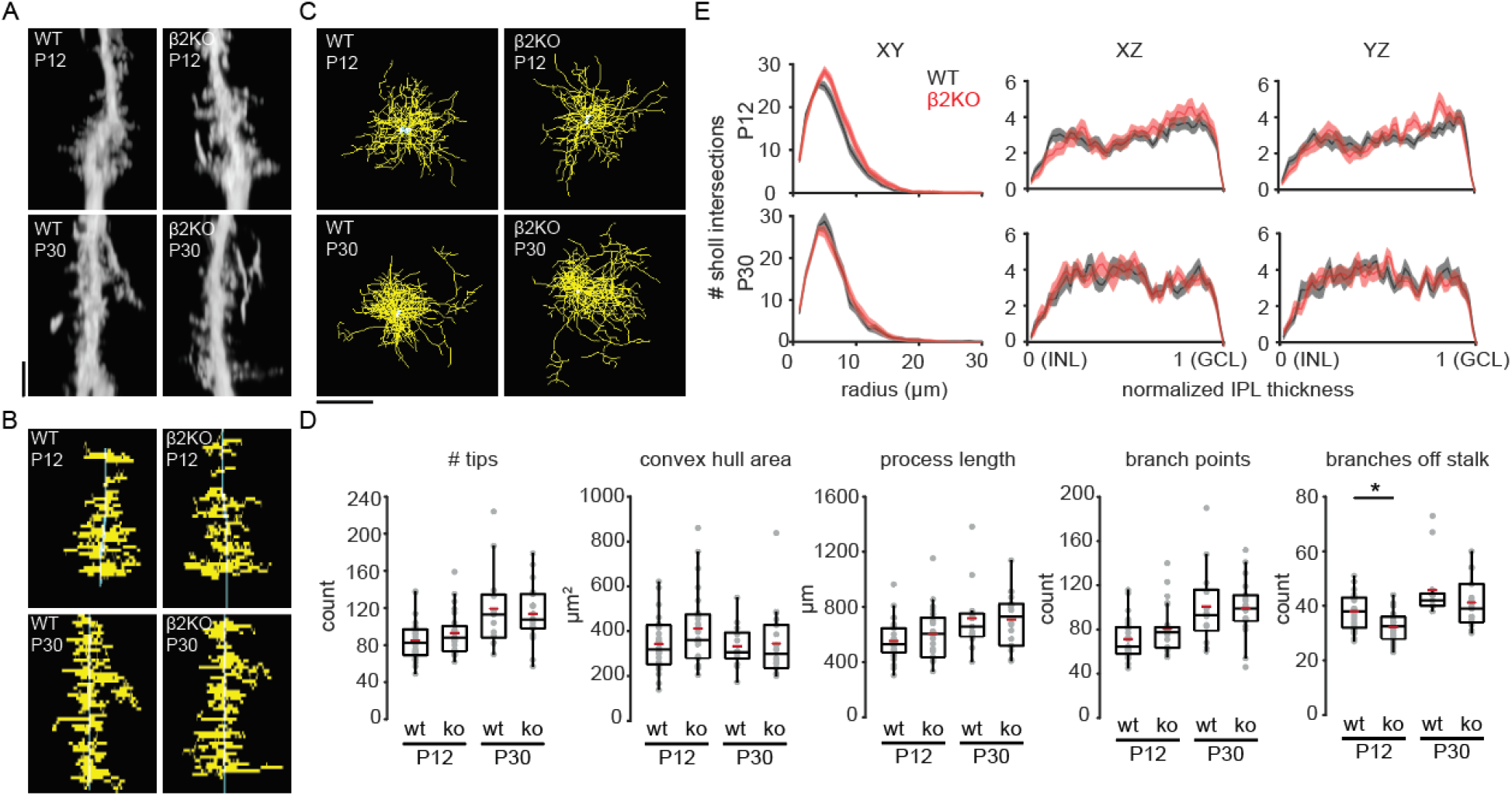
Chronic perturbation of retinal waves does not impact distribution, length, or complexity of Müller glial lateral. **(A)** Orthogonal projections of two-photon Z stacks obtained after sharp-filling single Müller glial cells with Alexa-488 at P12 (top) and at P30 (bottom) in wild type (WT, left) and in β2 nicotinic acetylcholine receptor knockout retinas (β2KO, right). Image scale bar is 10μm. **(B)** XZ projections of skeletons from cells shown in (A). **(C)** XY projections of skeletons from cells shown in (A). **(D)** Morphological measurements obtained from traced and skeletonized Müller glia. See Figure 6-figure supplement 1 for summary statistics. **(E)** Sholl intersection profiles for wild type (black) and β2KO (red) at P12 (top) and P30-39 (bottom). Left plots are Sholl profiles carried out on XY projections, middle plots are from XZ projections, and right plots are from YZ projections. For XZ and YZ projections, the center of the Sholl radius was placed on portion of the stalk closest to the inner nuclear layer (INL). GCL; ganglion cell layer. Source data available in Figure 6-source data 1.

## Discussion

In this study, we showed that as Müller glial lateral processes emerge during the second postnatal week of development, they exhibit both rapid motility and retinal wave-associated calcium transients that are partially compartmentalized. Global calcium transients in stalks were delayed relative to transients in lateral processes, were inhibited by an M1 mAChR antagonist, and became smaller and less frequent after P10. In contrast, wave-associated calcium transients in lateral processes often occurred locally and independent of stalk transients, were less sensitive to M1 mAChR block, and persisted throughout the second postnatal week. However, in contrast to astrocytes and radial glia in other regions of the central nervous system (CNS)^12,16,22,24^, Müller glia lateral process motility and outgrowth occurred independent of neuronal signaling and calcium transients. These results indicate that glial process morphology in the IPL is not regulated by neuronal activity.

### Mechanisms underlying Müller glial process motility

Glial morphology is critical for normal development of circuits. For example, a plexus of glial processes provides a diffusional barrier for neurotransmitters and other signaling molecules^7,41^, undergoes contact-mediated signaling with neurites to modulate synapse formation and function^42,43^, and participates in synaptic pruning via activation of phagocytic pathways in glia^44^.

Across brain regions, neuron-glia signaling impacts glial morphology in diverse ways. We found in the retina that glial lateral process motility during development was not impacted by neuron-glia signaling via release of neurotransmitter. This is similar to Bergmann glia, radial glia of the cerebellum, whose process motility is developmentally regulated and not directly affected by perturbations of neuronal activity or calcium influx^16^. Similarly in microglia, another highly motile cell type, chelation of calcium with BAPTA-AM slightly slowed but did not block process motility^45^. These findings stand in contrast to perisynaptic processes of hippocampal astrocytes, in which metabotropic glutamate receptor (mGluR) activation leads to localized increase in intracellular calcium which promotes process motility and coverage of dendritic spines^22^. This divergence between Müller glia and hippocampal astrocytes might be reflective of differences in ultrastructural interactions between glial processes and synapses in these two systems: perisynaptic processes of hippocampal astrocytes engage in true tripartite synapses in a brain region with high plasticity^46,47^, while there is currently no published evidence revealing similar structures in the retina. Further, astrocytic processes express specific actin-binding proteins such as ezrin which enable coupling of cytoskeletal dynamics with signaling through mGluRs^48^, which were previously shown to have minimal contribution to retinal wave-associated responses in Müller glia^20^.

Multiple pathways independent of neuronal activity may regulate morphological dynamics in Müller glial processes. One possibility is the release of growth factors from neurons or other nearby cells, which has been implicated in morphological changes among astrocytes in other systems^49,50^. This hypothesis is supported by our finding that application of exogenous EGF enhanced process motility prior to eye opening. Interestingly, EGFR expression in Müller glia is high prior to eye opening and declines with a developmental time course that matches our observed decline in motility^27^. Another intriguing possibility is that repulsive homotypic interactions between lateral processes from neighboring Müller glia underly their motility. This idea is supported by single-cell photoablation experiments in zebrafish Müller glia and mouse Schwann cells, in which processes from non-ablated cells filled in the territory vacated by ablated cells^51,52^. Further study will be required to determine whether these or other alternative pathways mediate Müller glial lateral process motility during development.

### Calcium compartments in Müller glia

We observed two distinct calcium compartments in Müller glia activated by retinal waves. Compartmentalization of calcium within Müller glial stalks was achieved via activation of glial M1 mAChRs, while lateral processes exhibited non-M1 mAChR-mediated transients with a smaller latency than those in stalks. We observed an age-dependent reduction in stalk participation in waves, while lateral processes continued to respond through P12. (Note, our observation that lateral processes continue to respond to retinal waves from P9 – P12 contrasts with a previous study that reported an overall reduction in glial responses to waves at these ages^20^. We attribute this difference to improved sensitivity of GCaMP6f over GCaMP3).The reduced stalk participation observed in older animals may be due to downregulation of M1 mAChRs prior to eye opening, or to a reduction in volume release of acetylcholine during the transition to glutamatergic waves53. Our observation that gabazine application during glutamatergic waves caused wave-associated transients to return in stalks suggests that the latter is true.

Subcellular calcium compartmentalization plays a variety of roles in neuron-glia signaling and tissue homeostasis^54^. Hippocampal astrocytic processes undergo at least two types of spatiotemporally distinct calcium transients in response to nearby synaptic activity, depending on the type of synaptic activity that occurs. This calcium signaling is thought to regulate synaptic transmission within the astrocytic territory^11^. Similarly, compartmentalized calcium transients during startle response frequently occur in distal branches, and less frequently in somata of hippocampal astrocytes in awake mice. Distinct signaling pathways, including those involving transmembrane calcium channels or transporters, adrenergic receptors, and IP3 receptors, differentially contribute to calcium transients in branches vs. somata^55^. Another study used a computational approach to reveal how calcium influx in fine astrocytic branches can be mediated by glutamate transporter-dependent activity of Na^+^/Ca^++^ exchangers, while somatic calcium can be modulated by mGluR-dependent activation of IP3 signaling^56^. Compartmentalization between Müller glial stalks and processes may arise by a similar mechanism.

We found that wave-associated compartmentalized calcium activity is not required for glial process motility, so this activity likely plays a role in other functions of Müller glia during retinal development. These functions include calcium-dependent neurovascular coupling^57^, release of gliotransmitters such as ATP or D-serine^17,18,58^, or secretion of synaptogenic molecules such as thrombospondin or growth factors^5,59^. Our identification of M1 mAChR as a driver of wave-associated calcium transients in stalks provides a target for selective perturbation in Müller glia in order to better define a role for these transients in retinal circuit development.

## Materials and methods

### Animals

All mice were purchased from The Jackson Laboratory and were maintained on mixed C57BL/6 backgrounds. For motility experiments, P8-P116 *GLAST-CreER;^mTmG^* mice were generated by cross breeding *GLAST-CreER* mice (strain 012586) with *Rosa26mTmG* mice (strain 007676)^14^. *GLAST-CreER* mice express tamoxifen-inducible Cre recombinase under control of a glia-specific promoter. *mTmG* is a dual-fluorescence reporter line which constitutively expresses membrane-bound tdTomato, and upon Cre-mediated recombination expresses membrane-bound green fluorescent protein (mGFP). For calcium imaging experiments, P9-P12 *GLAST-CreER;GCaMP6f* mice were generated by crossing *GLAST-CreER* mice with *GCaMP6f* (cytosolic) mice (strain 024105) or *Lck-GCaMP6f* (membrane-bound)^33^ mice (strain 029626) to enable specific and inducible calcium indicator expression in Müller glia. For sharp fills of Müller glial cells with fluorescent dye, we used C57BL/6 mice as controls for comparison with mice lacking the β2 subunit of the nicotinic acetylcholine receptor (β2-nAChR-KO).

Cre-mediated recombination was induced via intraperitoneal injection of 4-hydroxytamoxifen (50:50 E and Z isomers, Sigma-Aldrich) dissolved in sunflower seed oil. For uniform expression across the retina in neonates, injections of 0.5 mg tamoxifen were made 2 and 4 days before each experiment. For sparse expression to enable resolution of individual Müller glia during imaging, a single injection of 0.1 mg tamoxifen was made 2 days before each experiment.

All animal procedures were approved by the University of California, Berkeley Animal Care and Use Committee and conformed to the NIH Guide for the Care and Use of Laboratory Animals, the Public Health Service Policy, and the SFN Policy on the Use of Animals in Neuroscience Research.

### Retinal preparation

Animals of either sex were anesthetized via isoflurane inhalation and decapitated. Eyes were enucleated and retinas dissected in oxygenated (95% O_2_/5% CO2) artificial cerebrospinal fluid (ACSF) at room temperature under bright field (less than P10) or infrared (P10 and above) illumination. ACSF contained [in mM] 119.0 NaCl, 26.2 NaHCO3, 11 glucose, 2.5 KCl, 1.0 K2HPO4, 2.5 CaCl2, and 1.3 MgCl2. Isolated retinas were mounted ganglion cell side up on filter paper (Millipore) and transferred into the recording chamber of an upright microscope for imaging and electrophysiological recording. Retinas were continuously superfused with oxygenated ACSF (2-4 ml/min) at 32°-34°C for the duration of experiments and kept in the dark at room temperature in oxygenated ACSF when not imaging or recording. In a subset of experiments, wild type retinas were bath-loaded with the organic calcium dye Cal520 (12 μM) for 1.5-2 hours prior to performing calcium imaging.

### Two-photon imaging

Two-photon imaging of Müller glia in the IPL was performed using a modified movable objective microscope (MOM; Sutter Instruments) equipped with an Olympus 60X, 1.0 NA, LUMPlanFLN objective (Olympus America). Two-photon excitation was evoked with an ultrafast pulsed laser (Chameleon Ultra II; Coherent) tuned to 920 nm for all fluorophores. The microscope was controlled by ScanImage software (www.scanimage.org). Scan parameters were [pixels/line x lines/frame (frame rate in Hz)]: 256 x 256 (1.48–2.98), at 1–2 ms/line. When imaging GCaMP6f fluorescence in glial stalks and processes, the focal plane was set to ~1/3 the distance from the ganglion cell layer to the inner nuclear layer. Volumetric imaging of motility in mGFP-expressing glial processes and of surrounding tdTomato-expressing cells was achieved by acquiring sequential two-channel Z-stacks through the entire IPL, with slices 1 μm apart and averaging 4 frames per slice. Volumes were taken every 2 minutes for a total of 10 minutes of imaging lateral process dynamics.

### Electrophysiological recordings

Whole-cell voltage clamp recordings were made from whole mount retinas while simultaneously imaging GCaMP6f fluorescence. Under infrared illumination, RGC somas were targeted for voltage clamp recordings using glass microelectrodes with resistance of 3-5 MΩ (PC-10 pipette puller; Narishige) filled with an internal solution containing [in mM] 110 CsMeSO_4_, 2.8 NaCl, 20 HEPES, 4 EGTA, 5 TEA-Cl, 4 Mg-ATP, 0.3 Na_3_GTP, 10 Na_2_Phosphocreatine, and QX-Cl (pH 7.2 and 290 mOsm). The liquid junction potential correction for this solution was −10 mV. Signals were acquired using pCLAMP10 recording software and a MultiClamp 700A amplifier (Molecular Devices), sampled at 20 kHz and low-pass filtered at 2 kHz.

### Pharmacology

For pharmacology experiments, after 5-10 minutes of recording data in ACSF, pharmacological agents were added to the perfusion, and experimental recordings were obtained 5 minutes afterward. Drug concentrations were as follows: 5 μM cytochalasin-D (Avantor), 10 μM nocodazole (Sigma-Aldrich), 1 unit (100ng)/mL EGF (Cytoskeleton, Inc.), 5 μM pirenzepine (Tocris), 5 μM gabazine (Tocris), 25 μM DL-TBOA (Tocris), 20 μM DNQX (Tocris), 50 μM AP5 (Tocris), 200 μM BAPTA-AM (Tocris). DL-TBOA and BAPTA-AM were prepared in 0.1% DMSO. For calcium chelation experiments, whole mount retinas were incubated in BAPTA-AM or vehicle (ACSF/0.1% DMSO) for 1.5-2 hours, and then moved to ACSF for 30 minutes prior to imaging^60^. To verify that BAPTA-AM loading led to loss of retinal waves and calcium activity in Müller glia, we imaged GCaMP6f fluorescence in *GLAST-CreER;cyto-GCaMP6f* or *GLAST-CreER;Lck-GCaMP6f* retinas at P10 before and after BAPTA-AM loading.

### Focal agonist iontophoresis

For focal agonist application to mGFP-expressing lateral processes in *GLAST-CreER;mTmG* retinas, carbachol (10 mM, Tocris) or glutamic acid (10 mM, Sigma-Aldrich) was loaded into sharp electrodes pulled on a P-97 Micropipette Puller (Sutter) with a resistance of 100-150 MΩ. Electrodes also contained 2 mM Alexa-594 for verification of iontophoresis and to visualize electrodes under 2-photon illumination. While imaging at 920nm and visualizing fluorescence from mGFP and Alexa-594, electrodes were driven into the IPL at a ~15° angle using a micromanipulator set to the ‘DIAG’ function. For each cell, the tip of the electrode was placed 1-5 μm from the arbor of lateral processes of interest. Iontophoresis of agonist or control was achieved by applying continuous current using pCLAMP10 software. To apply carbachol, which is positively charged at physiological pH, continuous current of +4 nA was used, while −4 nA current was used to apply glutamate, which is negatively charged. Current of opposite polarity to that used for each respective agonist application was applied as a control, and the order of control vs. agonist application was shuffled to negate potential effects resulting from damage caused by the electrode. In a separate set of experiments, iontophoresis of agonist was performed in *GLAST-CreER;cyto-GCaMP6f* or *GLAST-CreER;Lck-GCaMP6f* retinas to verify that this method reliably evoked calcium transients in lateral processes.

### Glial sharp fills

To compare Müller glial morphology in wild type and β2-nAChR-KO retinas, sharp electrodes were pulled as described above, and the tip was bent 15-20° using a microforge. Electrodes were loaded with 2mM Alexa-488 in water, and iontophoresis of dye into single Müller glial cells was achieved by applying a −10 nA pulse for 500 ms in MultiClamp 700A software while the electrode was on the membrane of Müller glial endfeet in the ganglion cell layer. Electrodes were withdrawn as soon as cells started to fill with dye, and cells were imaged 1-5 minutes after filling^61^. 2-photon volumetric images of dye-filled lateral processes of single Müller glial cells in the IPL were acquired using the same image parameters used for imaging motility, as described above.

### Image processing and analysis: Calcium imaging

All images were processed using custom scripts in FIJI/ImageJ (National Institutes of Health)^62^ and MATLAB (MathWorks). For calcium imaging movies, following non-rigid motion correction^63^, regions of interest (ROIs) were semi-automatically placed over stalks using a filtering algorithm based on a Laplace operator and segmented by applying a user-defined threshold^64^. The Distance Transform Watershed function^65^ in FIJI was used to segregate nearby stalks in binarized images. This method defined most of the ROIs that an experienced user would recognize by eye. Manual adjustments were made to include stalks that were missed and to remove ROIs that were erroneously placed on structures other than stalks, which were defined as semiregularly spaced, punctate regions of fluorescence 2-3 μm in diameter in average intensity projection images. Lateral process ROIs were subsequently defined by randomly placing 250 2.5 μm x 2.5 μm squares in regions that did not overlap with stalk ROIs. Fluorescence intensity is reported as the average intensity across all pixels within the area of each ROI, and normalized as the relative change in fluorescence (ΔF/F) as follows:

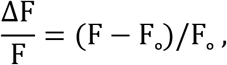

where F is the instantaneous fluorescence at any time point and F_o_ is the baseline fluorescence, defined as the median fluorescence value over the duration of the trace.

ΔF/F traces were smoothed using a 2-frame median filter and analyzed to detect calcium transients using custom MATLAB code. Traces were Z scored, and a threshold Z score of 3 was used to define calcium transients, which were then further defined as wave-associated if they occurred within 3 seconds of the peak of a wave-associated EPSC. Intercompartment latency was defined as the difference, in seconds, between wave-associated transients in stalks and the median wave-associated transient time for all lateral processes within the FOV, for each retinal wave.

### Image processing and analysis: Glial morphology

For volumetric images of glial process morphology, images were bandpass filtered in XY space to reduce noise while maintaining structure of fine processes, and registered using the Correct 3D Drift plugin in FIJI^66^. 10-minute volumetric time series were corrected for photobleaching using the ‘Histogram matching’ setting within FIJI’s Bleach Correction plugin. Time series were collapsed into temporally color-coded images to facilitate identification of motile and stable processes in 3D. The Cell Counter tool within FIJI was used to count lateral processes and define their morphological dynamics as one of the following: extending, retracting, new process, lost process, extension followed by retraction, retraction followed by extension, and stable. Locations of process tips within the IPL were recorded and binned into one of five equally sized sections, corresponding to putative sublayers S1 though S5, defined by their distance from the INL and GCL borders as identified using membrane-tdTomato fluorescence on somas.

Volumetric images of sharp-filled Müller glia were traced using the Simple Neurite Tracer plugin (FIJI)^67^. We traced each glial stalk and subsequently any visible processes branching from the stalk, creating distinct paths for each process and preserving branch order relationships. Measurements including number of tips, total process length, total branch points, and primary branches from stalk were derived from traced paths. For Sholl and convex hull analyses, paths were converted to binary skeletons and registered to correct for XY displacement of the stalk between the GCL and INL. Sholl analysis was performed using concentric rings spaced 1 μm apart^68^. For the XY plane, the center of each Sholl radius was placed on the registered stalk, and intersections were counted at each radius overlayed on a maximum projection image. For the XZ and YZ planes, the center of each Sholl radius was placed on the end of the stalk closest to the INL in orthogonal projections of traced cells. Sholl radii were normalized to the total IPL thickness. Convex hull area was defined as the area of the smallest convex polygon enclosing the entire skeleton in an XY maximum projection image.

### Statistical analysis

Group measurements are expressed as mean ± SEM. Chi-squared tests of goodness-of-fit were used to test for nonuniformity in lateral process distribution across the IPL within each age. Chi-squared tests for independent proportions were used to test for age-dependent differences in process distribution across the IPL (Figure 1-figure supplements 1 and 2). When comparing proportion of total stable processes between control and experimental manipulations, we applied Wilcoxon signed rank tests when cells were paired between conditions, and Wilcoxon rank sum tests for unpaired cells. When there were less than 5 cells in a particular condition, we pooled process counts between cells and report chi-square test statistics for these comparisons as well (Figure 1-figure supplements 3–5; Figure 5-figure supplement 1). To test for age-, compartment-, and drug-dependent differences in retinal wave-associated calcium transients (Figure 2-figure supplement 2; Figure 3-figure supplements 1–4; Figure 4-figure supplements 1–4), we applied two- or three-way mixed ANOVAs followed by post hoc t-tests with Benjamini-Hochberg correction for multiple comparisons, when appropriate. Paired t-tests were used to compare fold-changes between compartments in response to pirenzepine and gabazine (Figure 3-figure supplement 2 and 4; Figure 4-figure supplement 2 and 4). Intercompartment latency of wave-associated calcium transients was compared between ages and conditions using Wilcoxon rank sum tests, and within each condition tested for significant difference from zero using Wilcoxon signed rank tests (Figure 2-figure supplement 3; Figure 3-figure supplement 5; Figure 4-figure supplement 5). Wave EPSC amplitude and IWI were compared between control and drug conditions using paired t-tests (Figure 3-figure supplement 6; Figure 4-figure supplement 6). Sholl intersection profiles between wild type and β2-nAChR-KO Müller glia were compared using two-way repeated measures mixed ANOVA to test for genotype-associated differences in complexity across Sholl radii (Figure 6). Two-sample t-tests were used for comparison of other morphological measurements between wild type and β2-nAChR-KO Müller glia (Figure 6-figure supplement 1).

## Acknowledgements

We thank members of the Feller lab for commenting on the manuscript. JT was supported by the National Science Foundation Graduate Research Fellowship (DGE 1752814). J.T. and M.B.F. were supported by NIH grants R01EY019498, R01EY013528, and P30EY003176.

## Figure supplements

**Figure 1-figure supplement 1.**
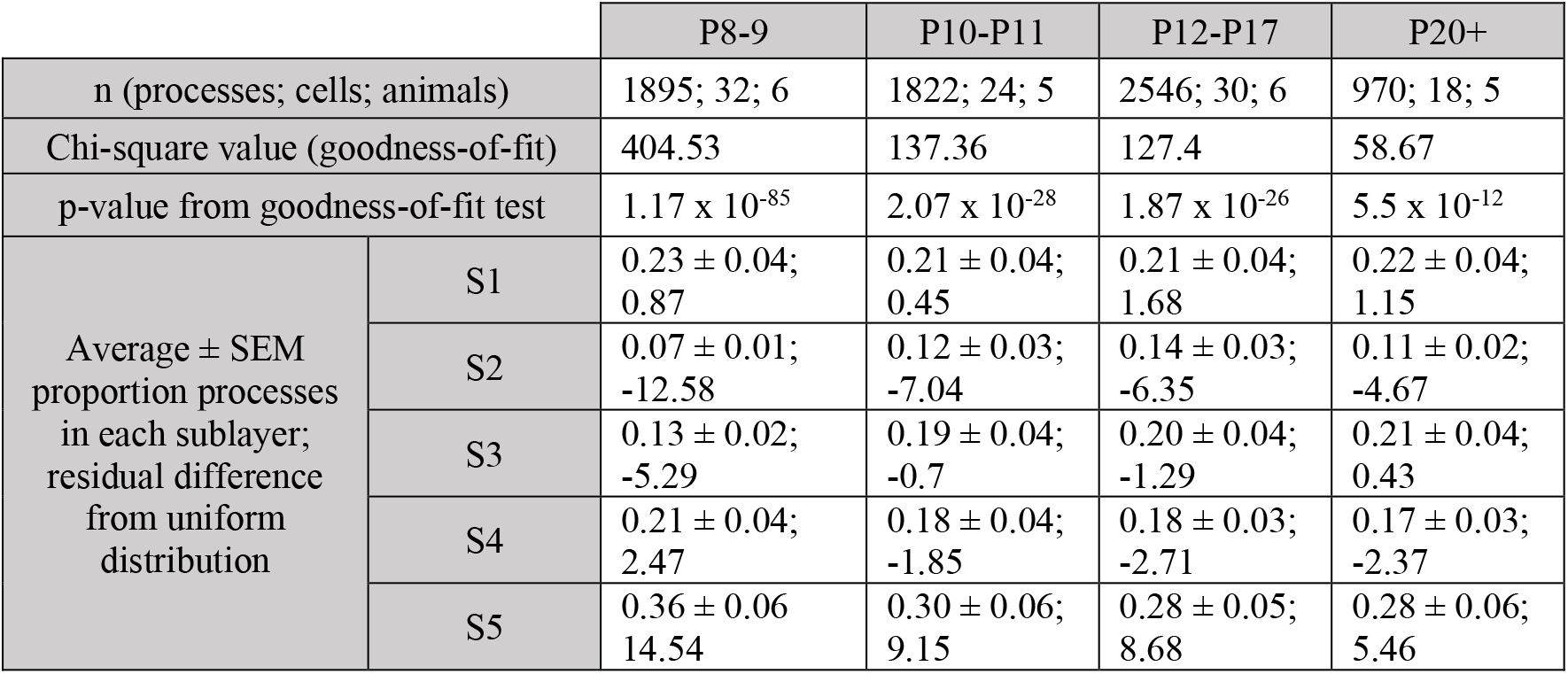
Analysis of differences in lateral process outgrowth between sublayers; separate tests for each age group Chi-square goodness-of-fit tests were used to test for nonuniform distribution of processes across the IPL at each age. p-values were corrected using Benjamini-Hochberg adjustment. Related to Figure 1D. Source data available in Figure 1-source data 1.

**Figure 1-figure supplement 2.**
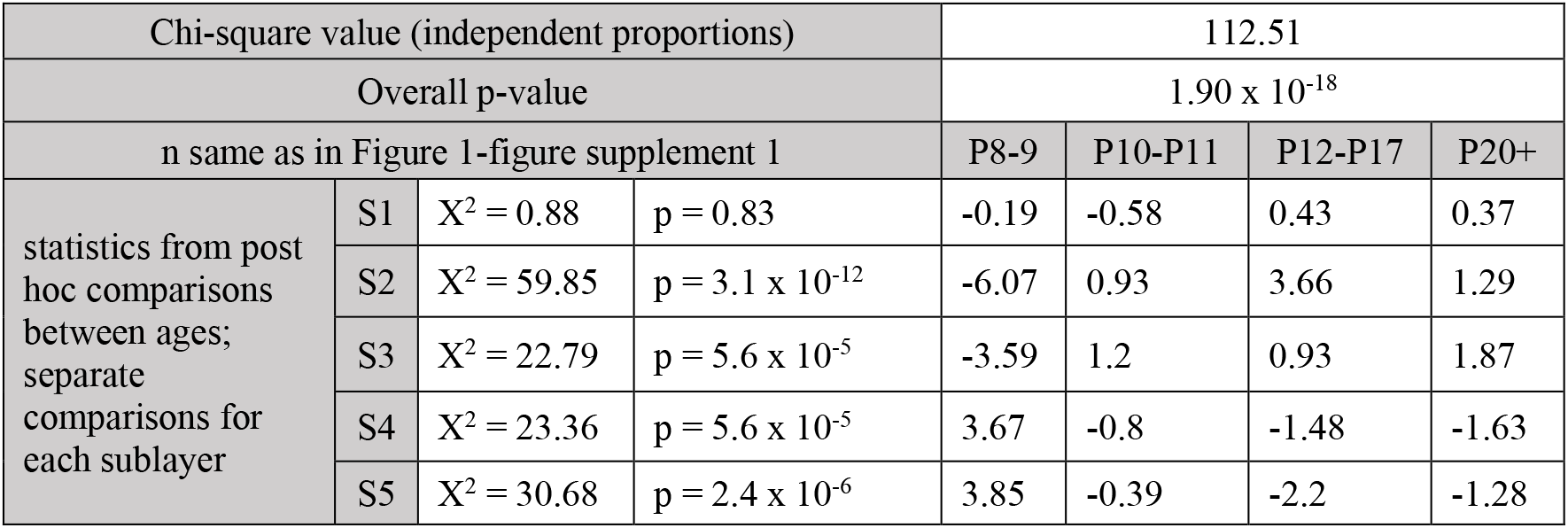
Analysis of age-dependent differences in lateral process outgrowth within each sublayer Following an overall Chi-square test for independent proportions across ages, separate Chi-square tests for independent proportions were performed for each sublayer to test for sublayer-specific developmental changes in process outgrowth. p-values were corrected using Benjamini-Hochberg adjustment. Residual differences in proportions across ages are reported for each Chi-square test. Related to Fig. 1D. Source data available in Figure 1-source data 1.

**Figure 1-figure supplement 3.**
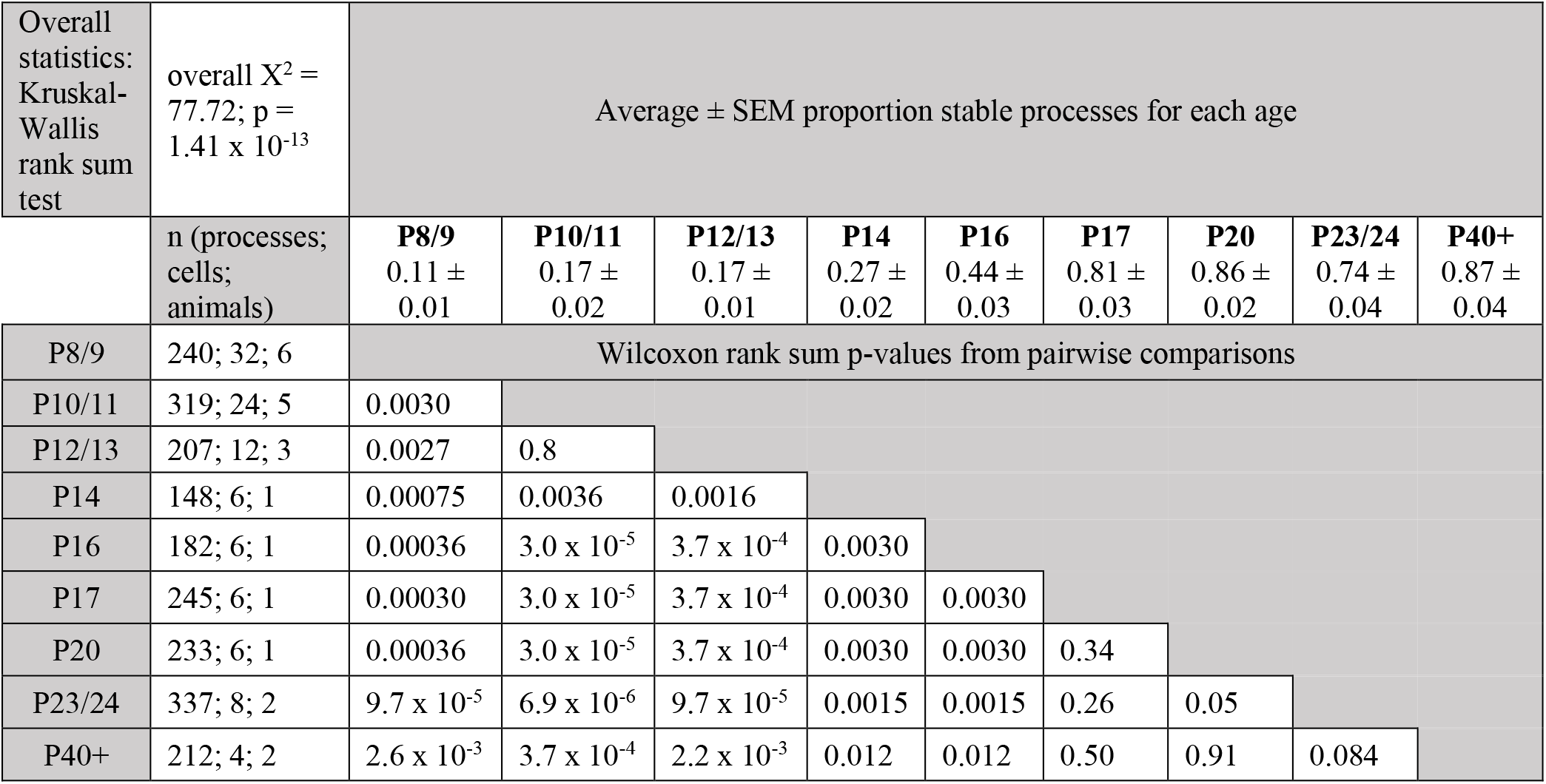
Analysis of age-dependent differences in proportion stable processes Following overall Kruskal Wallis rank sum test for age-dependent changes in proportion of total stable processes, separate pairwise Wilcoxon rank sum tests were performed to determine developmental time course of process stabilization. Reported p-values are false discovery rate-corrected. Related to Fig. 1F. Source data available in Figure 1-source data 1.

**Figure 1-figure supplement 4.**
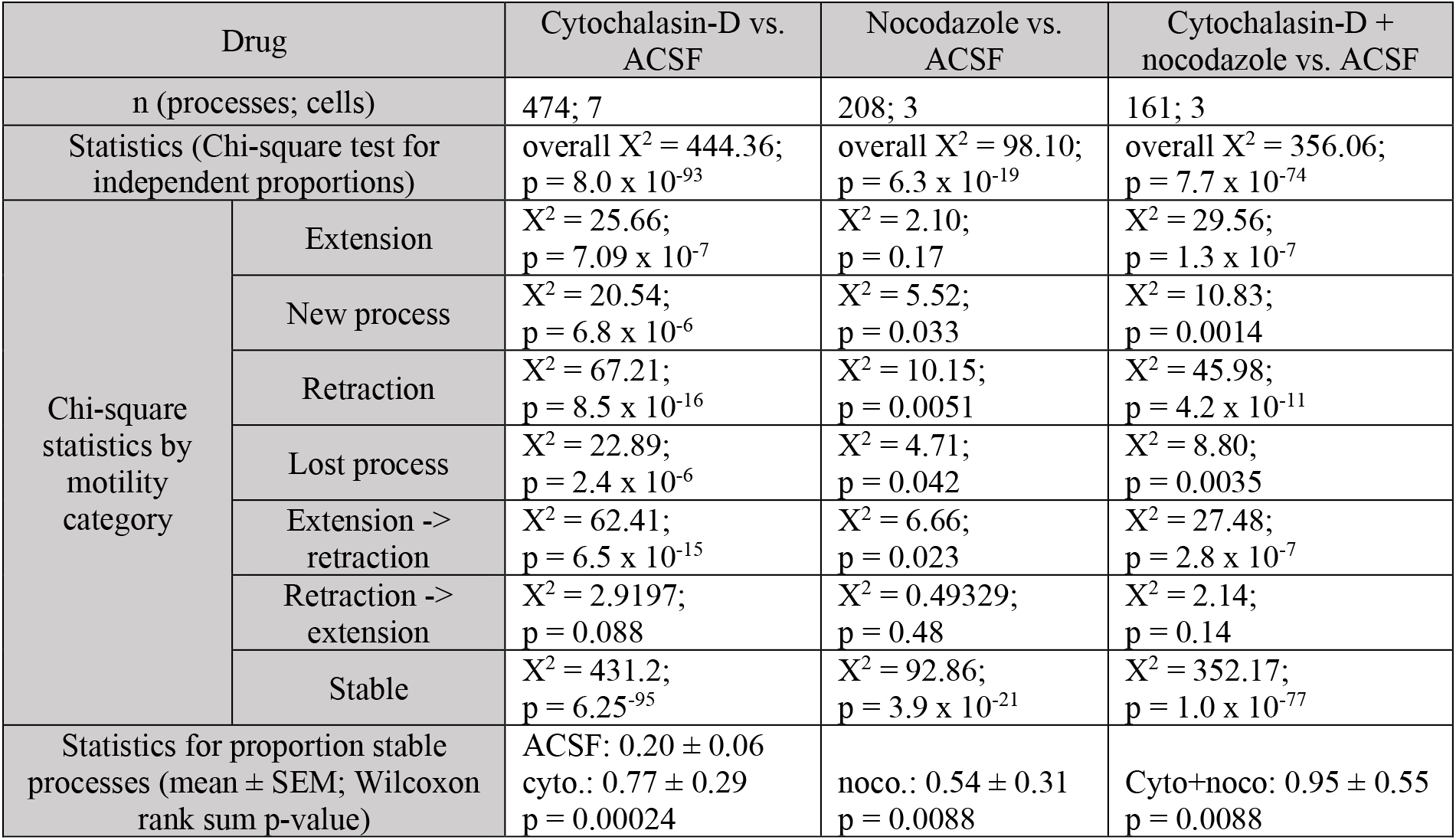
Analysis of motility in blockers of cytoskeletal rearrangements Chi-square tests for independent proportions were used to test for changes in motility in presence of Cytochalasin-D and nocodazole. Controls were in ACSF (n = 967 processes; 12 cells). Post hoc Chi-square tests were used to test for differences in proportions of processes undergoing each type of motility in control vs. drug. Pairwise comparison of proportion stable processes was performed using Wilcoxon rank sum test. p-values were corrected using Benjamini-Hochberg adjustment. Related to Fig. 1G. Source data available in Figure 1-source data 1.

**Figure 1-figure supplement 5.**
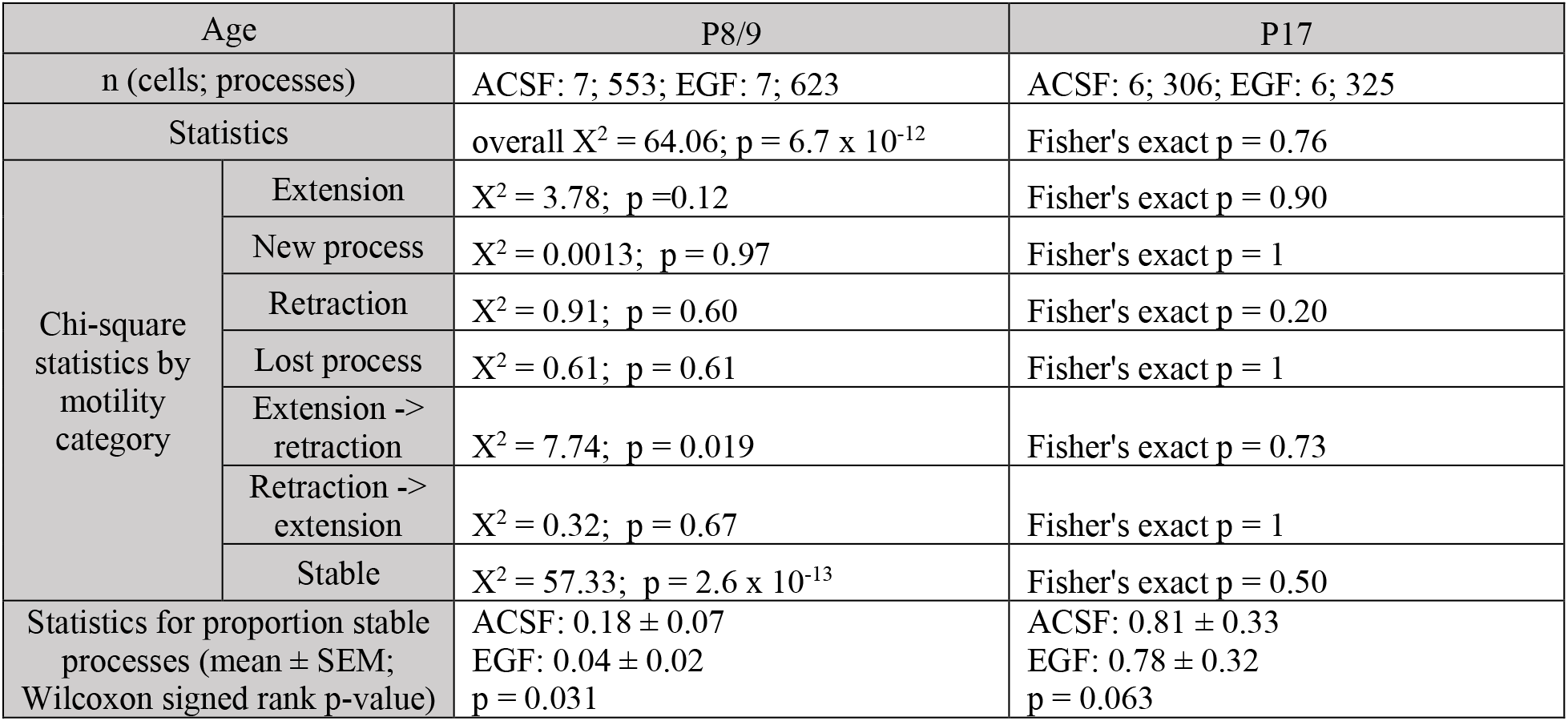
Analysis of motility in exogenous EGF Chi-square tests for independent proportions were used to test for changes in overall process motility in the presence of exogenous EGF. Post hoc Chi-square tests were used to test for differences in proportions of processes undergoing each type of motility in control vs. EGF. Pairwise comparison of proportion stable processes was performed using Wilcoxon signed rank test. p-values were corrected using Benjamini-Hochberg adjustment for multiple comparisons. Related to Fig. 1H. Source data available in Figure 1-source data 1.

**Figure 2-figure supplement 1.**
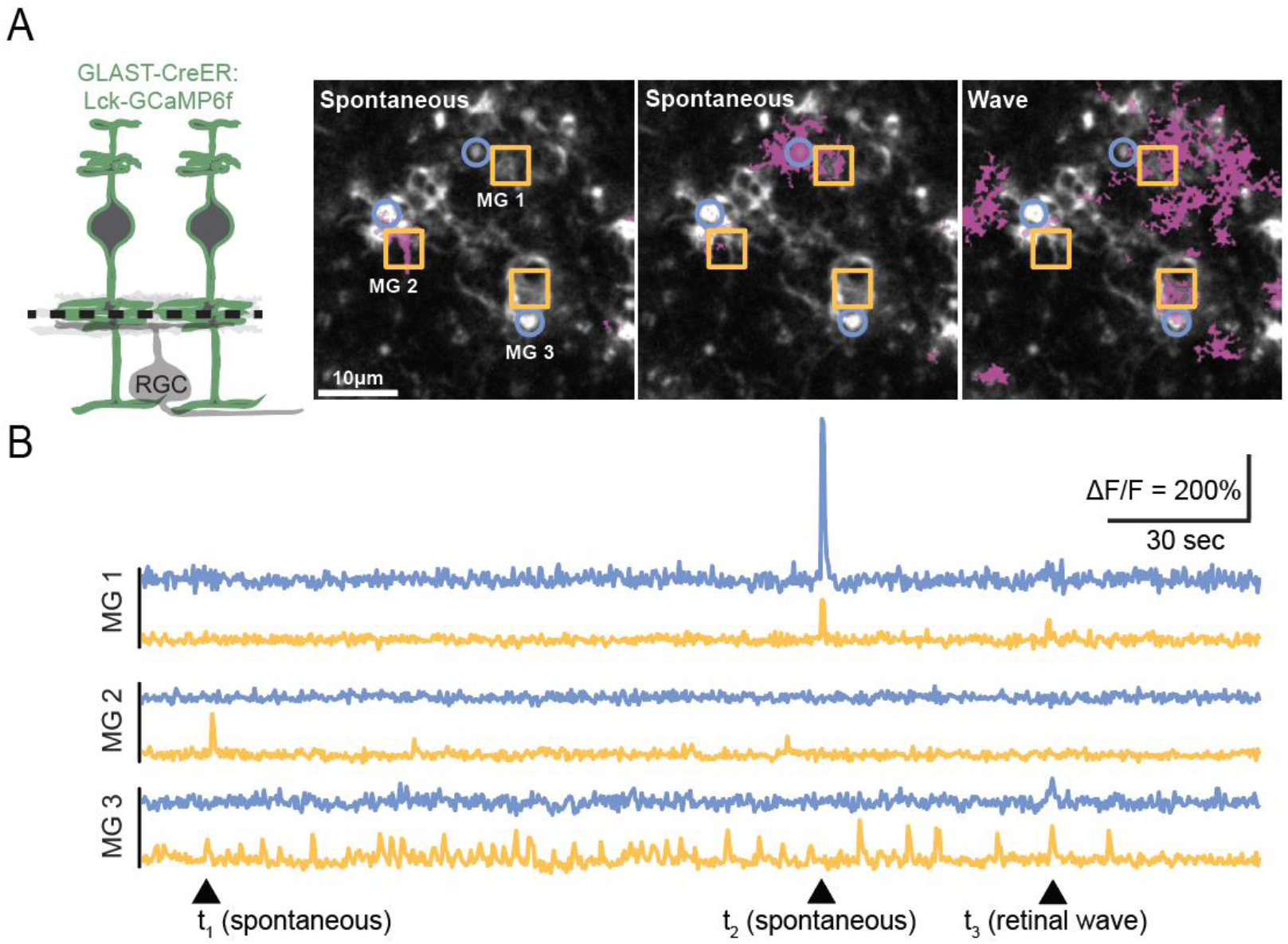
Visualization of calcium compartments using *GLAST-CreER;Lck-GCaMP6f*. **(A)** *Left,* diagram of *GLAST-CreER;Lck-GCaMP6f* retina used for imaging calcium in glial processes. Black dashed line indicates 2-photon imaging focal plane. *Right,* example FOV showing two spontaneous calcium events (left and middle; transients denoted as magenta regions) and one wave-associated calcium event (right) in Müller glia (MG) at P10. Three stalks (blue circles) and three neighboring lateral process ROIs (yellow squares) have been denoted as MG 1-3. **(B)** Normalized fluorescence traces showing calcium activity in MG 1-3. Arrowheads indicate time points shown in images in (A).

**Figure 2-figure supplement 2.**
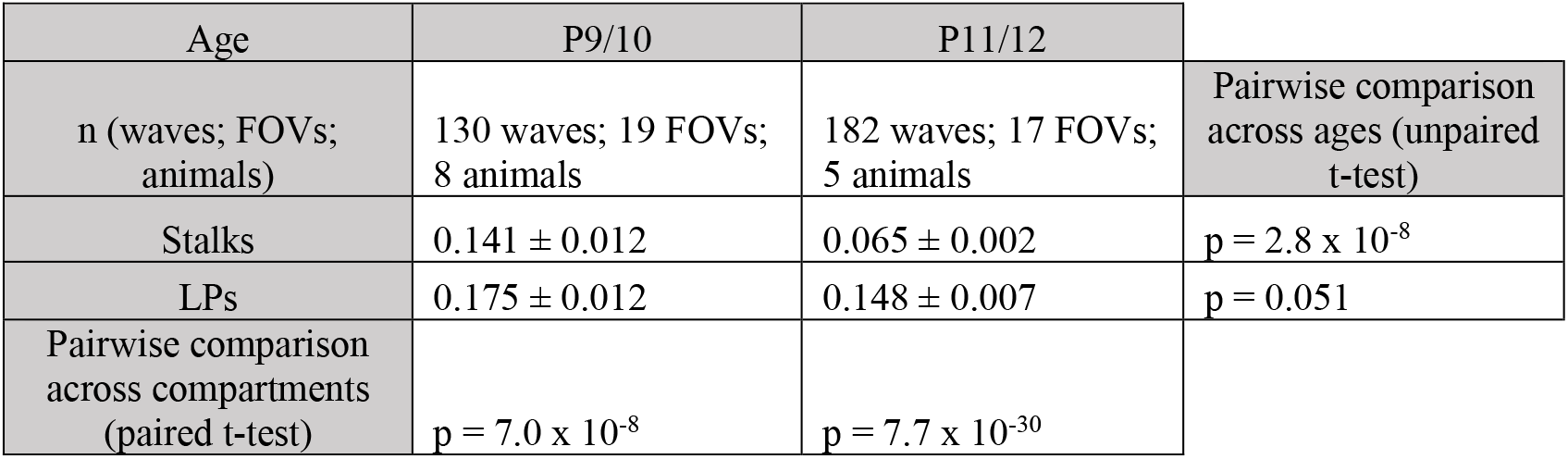
Analysis of calcium compartmentalization: proportion of ROIs participating in waves Two-way mixed ANOVA (within group: compartment) revealed significant main effects of age (p = 7.2 x 10^−6^) and compartment (p = 9.6 x 10^−33^), with a significant interaction between these factors (p = 7.6 x 10^−8^). Unpaired t-tests were used to compare proportion ROIs responding between ages, and paired t-tests were used to compare proportion responding between stalks and LPs within ages. Related to Fig. 2C. Source data available in Figure 2-source data 1.

**Figure 2-figure supplement 3.**
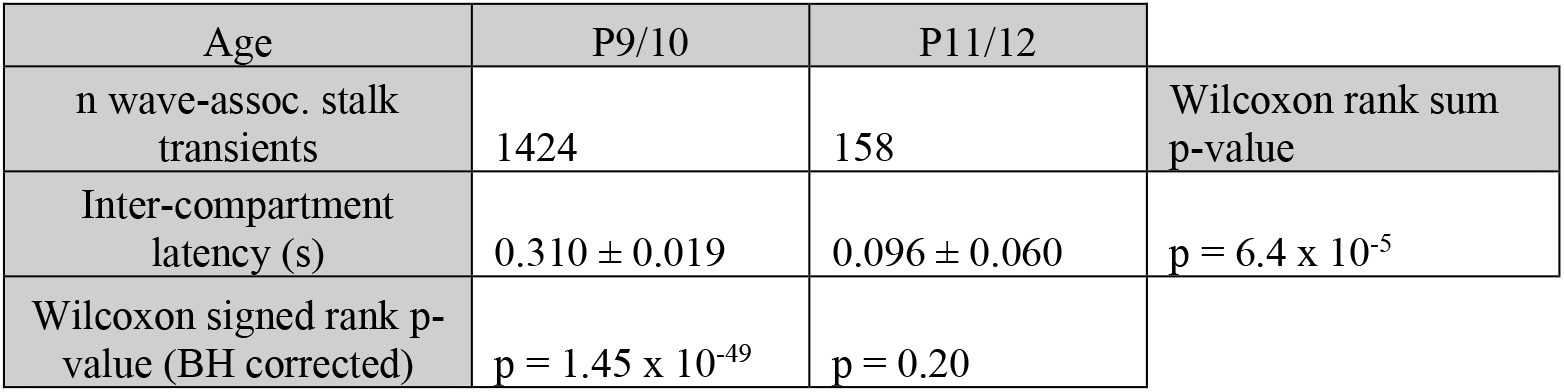
Analysis of intercompartment latency in ACSF Inter-compartment latency was calculated by subtracting the median wave-associated transient time among LPs during a retinal wave from individual wave-associated transient times among stalks responding to that wave. Wilcoxon signed rank tests were used to test for a non-zero latency at each age, and rank sum test was used to test for an age-dependent change in latency. Related to Fig. 2D. Source data available in Figure 2-source data 1.

**Figure 3-figure supplement 1.**
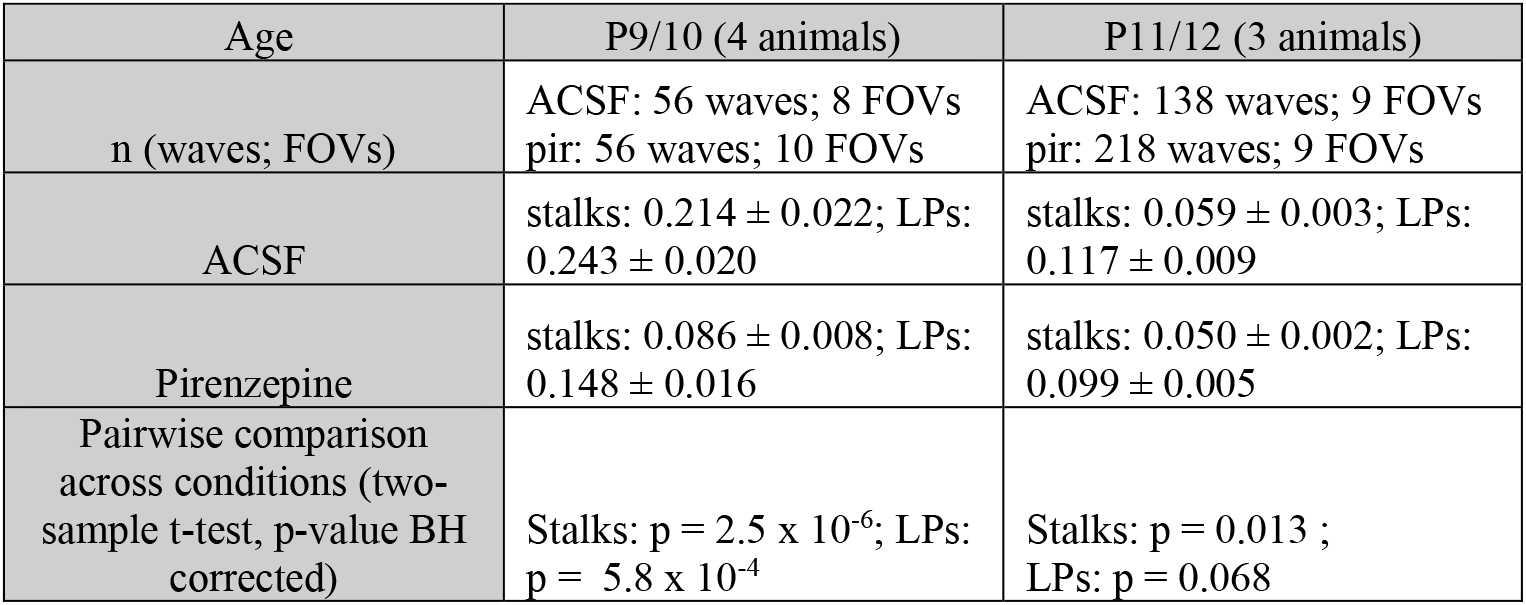
Proportion of total ROIs participating in retinal waves: ACSF vs. pirenzepine Three-way mixed ANOVA (within group: compartment) revealed significant main effects of drug (p = 2.7 x 10^−13^), age (p = 2.7 x 10^−13^), and compartment (p = 2.7 x 10^−13^) on proportion ROIs participating in waves, with a significant interaction between drug condition and age (p = 5.88 x 10^−9^). Two-sample t-tests were used for post hoc analysis of pirenzepine effect on proportion ROIs participating within stalks and LPs. Related to Fig. 3B, *middle.* Source data available in Figure 3-source data 1.

**Figure 3-figure supplement 2.**
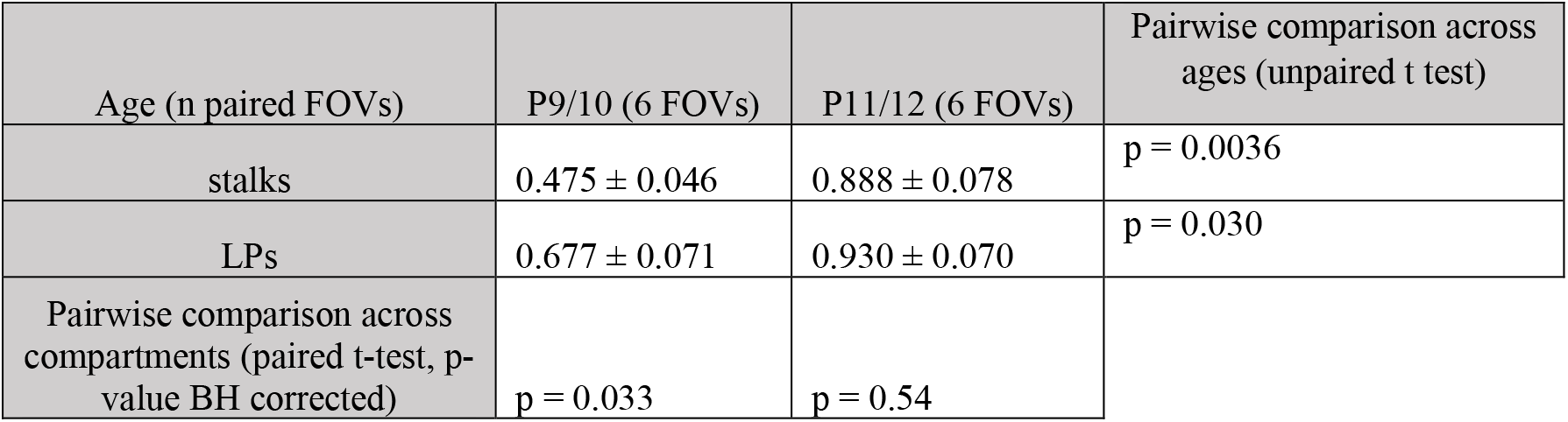
Fold change in proportion ROIs participating in retinal waves: ACSF vs. pirenzepine Two-way mixed ANOVA (within group: compartment) revealed significant main effects of age (p = 0.003) and compartment (p = 0.017) on fold change in proportion ROIs participating in waves during pirenzepine application. Paired t-tests were used for post hoc comparison of pirenzepine effect between stalks and LPs, and unpaired t-tests were used to compare pirenzepine effect between ages. Related to Fig. 3B, *right.* Source data available in Figure 3-source data 1.

**Figure 3-figure supplement 3.**
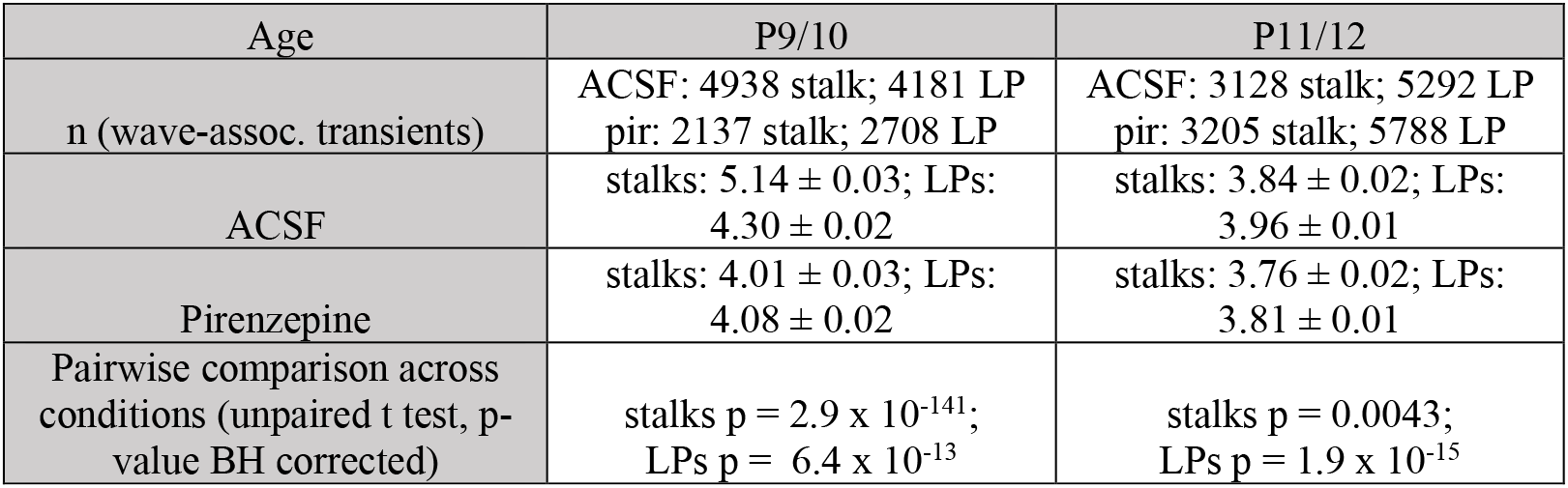
Z scored wave-associated ΔF/F amplitude: ACSF vs. pirenzepine Three-way repeated measures ANOVA revealed significant main effects of drug (p = 2.3 x 10^−127^), age (p = 2.1 x 10^−20^), and compartment (p = 1.7 x 10^−234^) on wave-associated transient amplitude, with significant interactions between drug:age (p = 2.2 x 10^−37^), drug:compartment (p = 6.6 x 10^−65^), and age:compartment (p = 1.4 x 10^−47^). Two-sample t-tests were used for post hoc analysis of pirenzepine effect on wave-associated transient amplitude within stalks and LPs. Related to Fig. 3C, *middle.* Source data available in Figure 3-source data 1.

**Figure 3-figure supplement 4.**
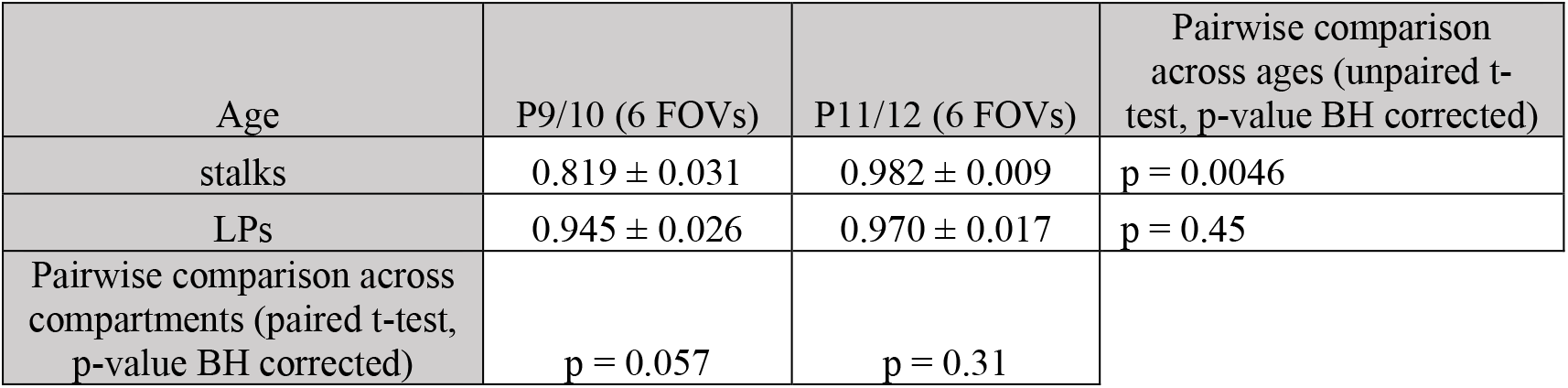
Fold change in wave-associated ΔF/F amplitude: ACSF vs. pirenzepine Two-way mixed ANOVA (within group: compartment) revealed significant main effects of age (p = 0.003) and compartment (p = 0.023) on fold change in wave-associated transient amplitude during pirenzepine application, with a significant interaction between age and compartment (p = 0.009). Paired t-tests were used for post hoc comparison of pirenzepine effect between stalks and LPs, and unpaired t-tests were used to compare pirenzepine effect between ages. Related to Fig. 3C, *right.* Source data available in Figure 3-source data 1.

**Figure 3-figure supplement 5.**
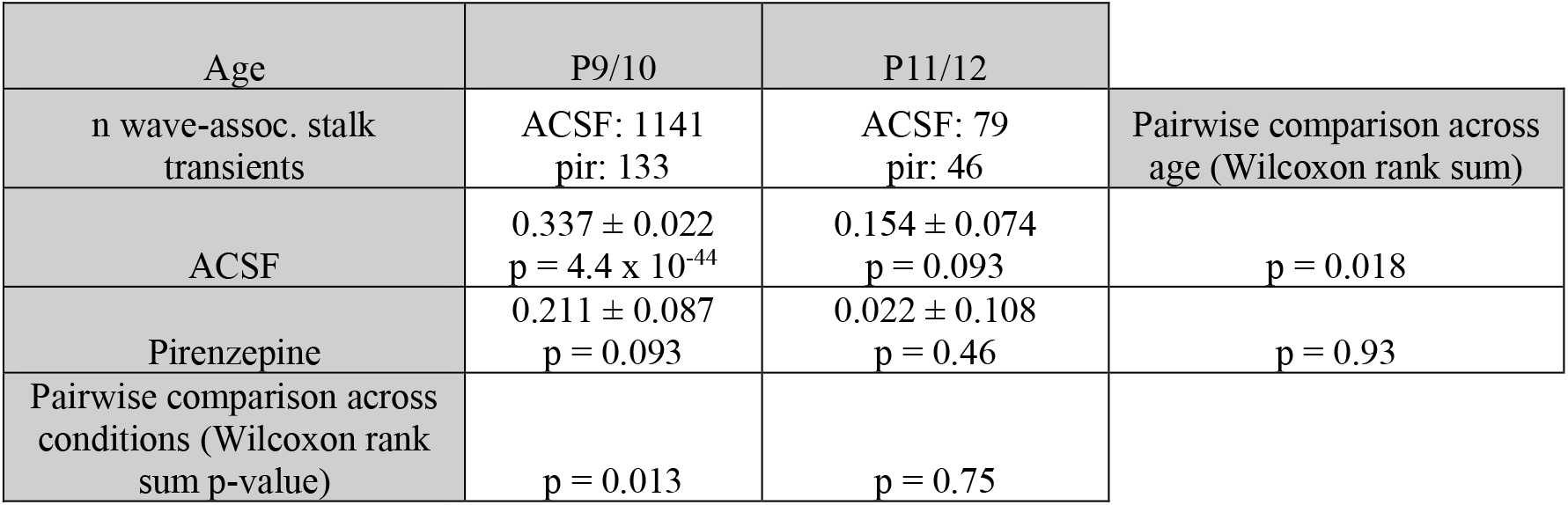
Intercompartment latency in ACSF vs. pirenzepine Wilcoxon signed rank tests were used to test for a non-zero latency within each condition and age, and rank sum tests were used to test for effects of pirenzepine and age on latency. Related to Fig. 3D. Source data available in Figure 3-source data 1.

**Figure 3-figure supplement 6.**
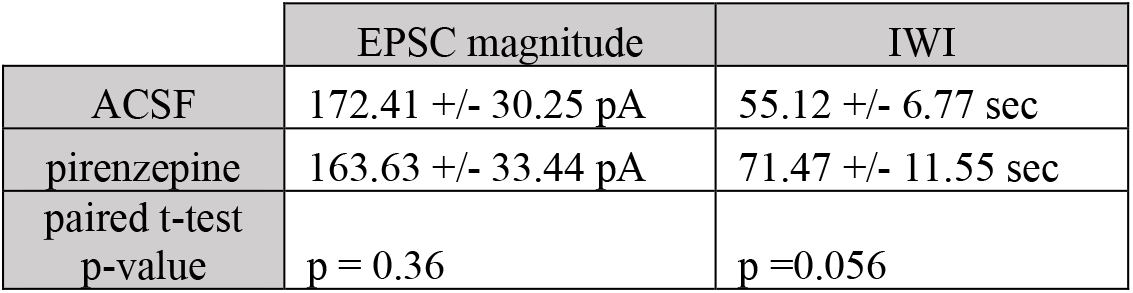
Wave properties in ACSF vs. pirenzepine Wave properties are averages per RGC voltage clamp recording (n = 13 cells). Wave-associated EPSC and inter-wave interval (IWI) were compared using paired t-tests. Related to Fig. 3E. Source data available in Figure 3-source data 1.

**Figure 3-figure supplement 7.**
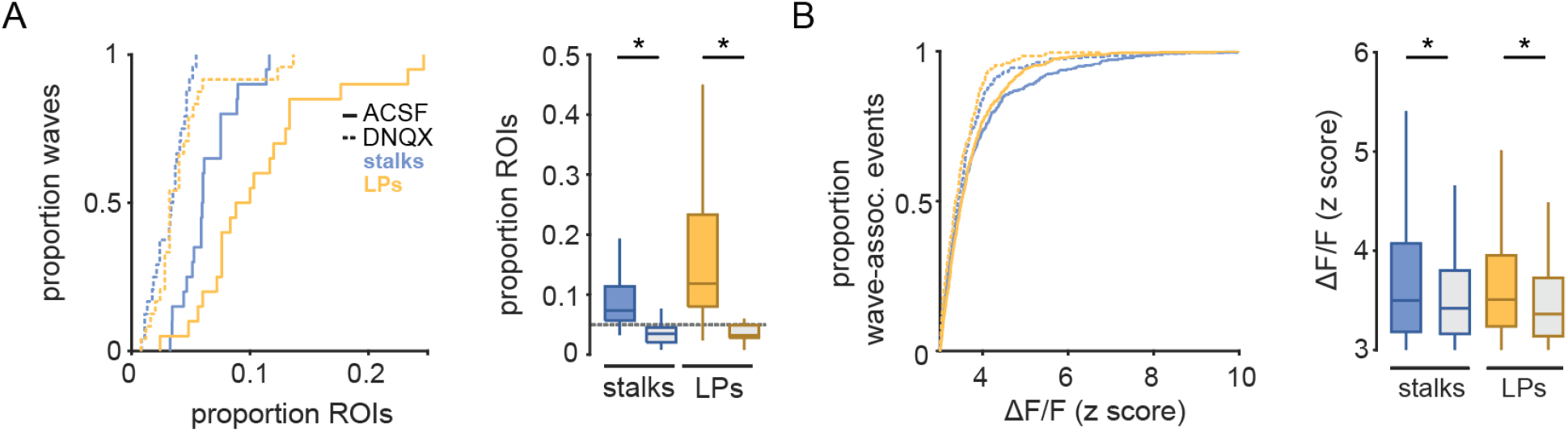
DNQX reduces glial participation in waves among stalks and lateral processes. **(A)** *Left,* cumulative distribution of proportion stalk ROIs (blue) or lateral process ROIs (yellow) undergoing wave-associated calcium transients for all waves in control (solid lines) and 20 μM DNQX (dotted lines). *Right,* box plots showing median and 25%-75% interquartile range for data shown in *left*. Colored boxes indicate control, while grey boxes indicate DNQX. Grey dashed line indicates proportion of ROIs undergoing calcium transients at randomly selected times. **(B)** *Left,* cumulative distribution of Z scored wave-associated ΔF/F amplitude in stalks and lateral processes in control and DNQX. *Right,* box plots showing median and 25%-75% interquartile range for data shown in *left.*

**Figure 4-figure supplement 1.**
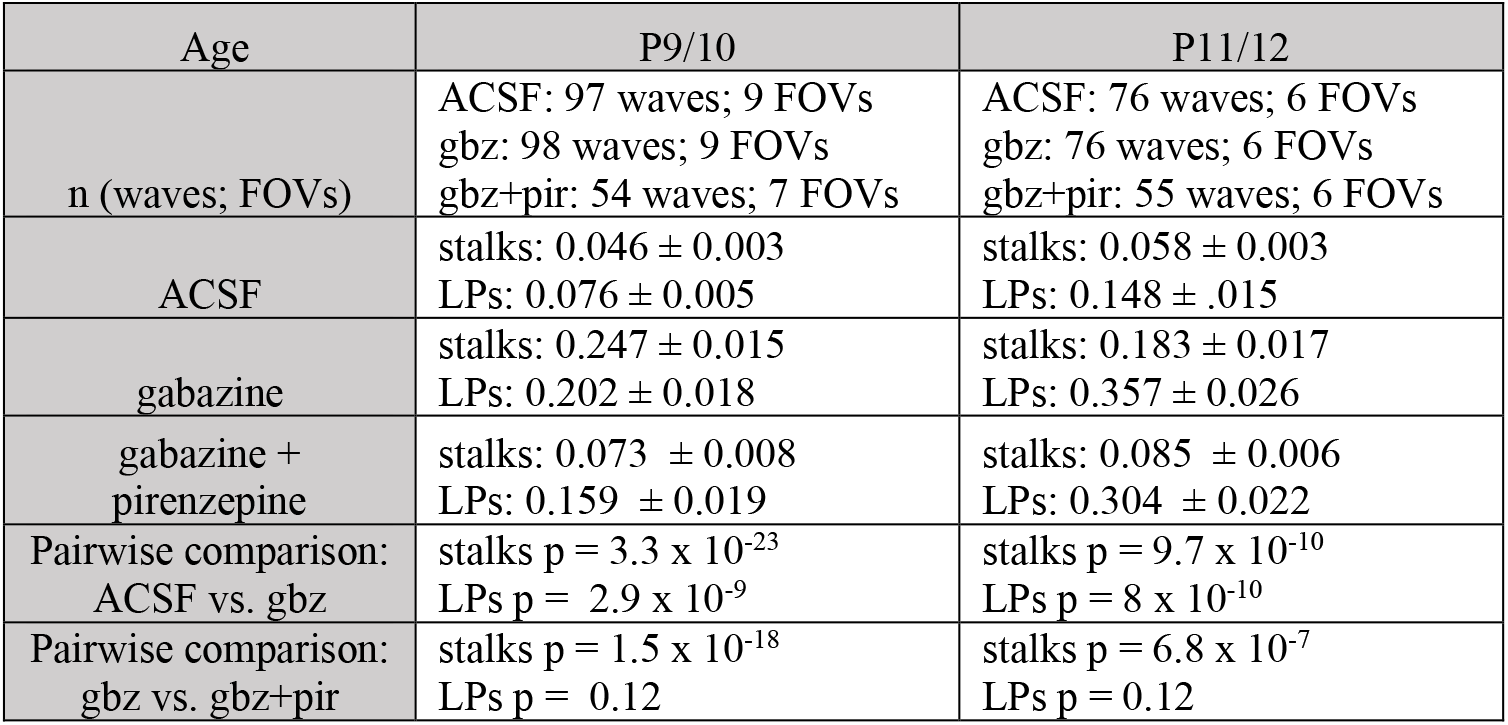
Proportion ROIs participating in retinal waves in ACSF, gabazine, and gabazine + pirenzepine Three-way mixed ANOVA (within group: compartment) revealed significant main effects of drug (p = 6.55 x 10^−32^), age (p = 1.4 x 10^−6^), and compartment (p = 1.8 x 10^−50^) on proportion ROIs participating in waves, with significant interactions between drug:compartment (p = 9.59 x 10^−12^), and age:compartment (p = 1.6 x 10^−31^). Two-sample t-tests were used for post hoc analysis of gabazine and pirenzepine effect on proportion ROIs participating within stalks and LPs. Related to Fig. 4B, *left.* Source data available in Figure 4-source data 1.

**Figure 4-figure supplement 2.**
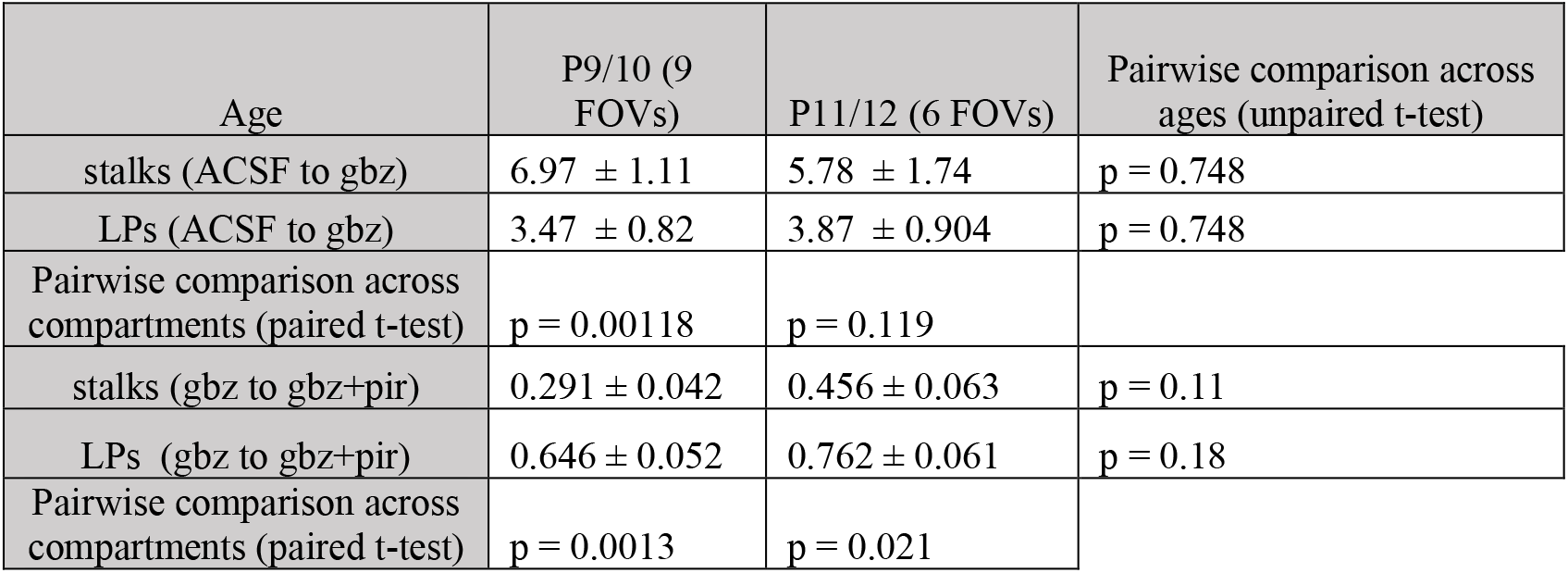
Fold change in proportion ROIs participating in waves in ACSF vs. gabazine, and in gabazine vs. gabazine + pirenzepine Two-way mixed ANOVA (within group: compartment) revealed significant effects of compartment (p = 0.00037 for ACSF to gabazine; p = 5.12 x 10^−5^ for gabazine to gabazine + pirenzepine), and of age (p = 0.03 for gabazine to gabazine + pirenzepine) on fold change in proportion ROIs participating in waves. Paired t-tests were used for post hoc comparison of gabazine and pirenzepine effect between stalks and LPs, and unpaired t-tests were used to compare drug effects between ages. Related to Fig. 4B, *right.* Source data available in Figure 4-source data 1.

**Figure 4-figure supplement 3.**
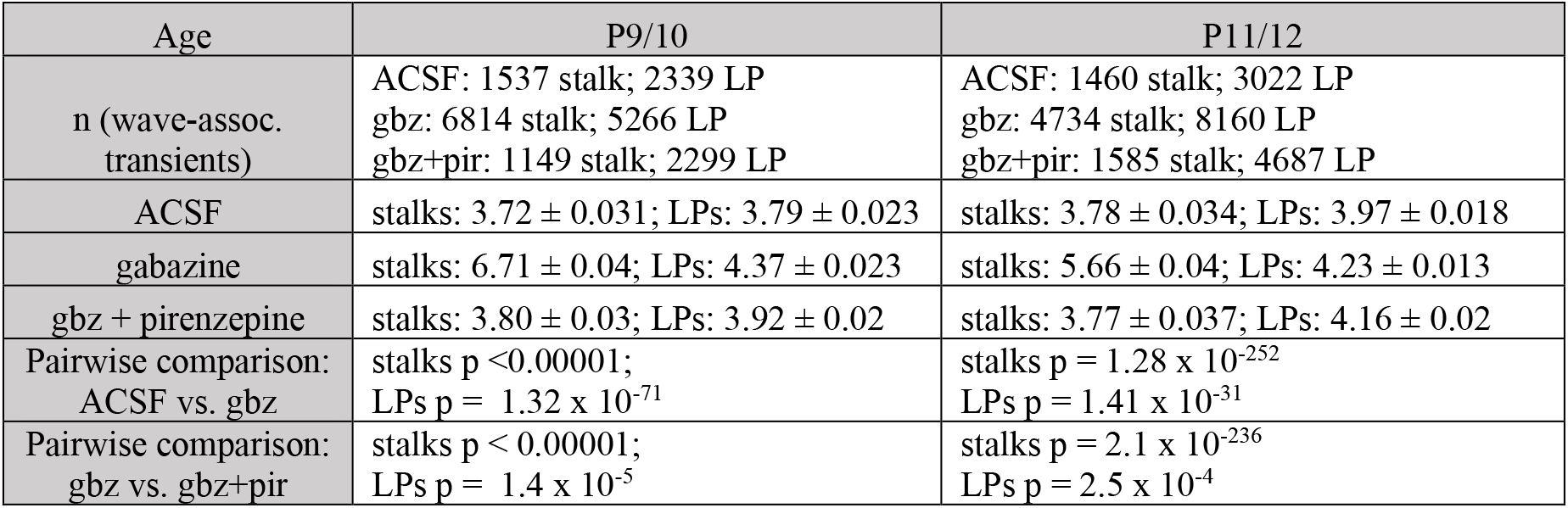
Z scored wave-associated ΔF/F amplitude in ACSF, gabazine, and gabazine + pirenzepine Three-way repeated measures ANOVA revealed significant main effects of drug (p < 0.00001), age (p = 2.2 x 10^−109^), and compartment (p = 7.3 x 10^−8^) on wave-associated transient amplitude, with significant interactions between drug:age (p < 0.00001), drug:compartment (p = 3.0 x 10^−67^), and age:compartment (p = 2.5 x 10^−22^). Two-sample t-tests were used for post hoc analysis of gabazine and pirenzepine effect on wave-associated transient amplitude within stalks and LPs. Related to Fig. 4C, *left.* Source data available in Figure 4-source data 1.

**Figure 4-figure supplement 4.**
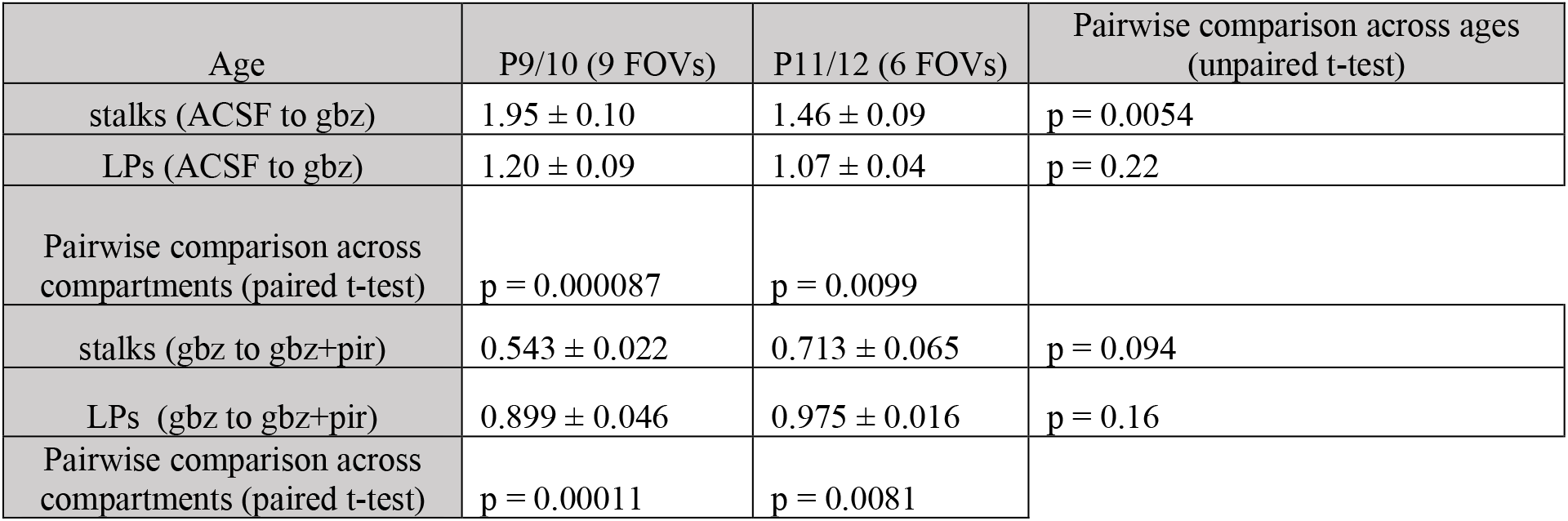
Fold change in wave-associated transient amplitude: ACSF vs. gabazine; gabazine vs. gabazine + pirenzepine Two-way mixed ANOVA (within group: compartment) revealed significant main effects of age (p = 0.014) and compartment (p = 1.8 x 10^−6^) on fold change in wave-associated transient amplitude during gabazine and pirenzepine application, with a significant interaction between age and compartment (p = 0.024). Paired t-tests were used for post hoc comparison of gabazine and pirenzepine effects between stalks and LPs, and unpaired t-tests were used to compare drug effects between ages. Related to Fig. 4C, *right.* Source data available in Figure 4-source data 1.

**Figure 4-figure supplement 5.**
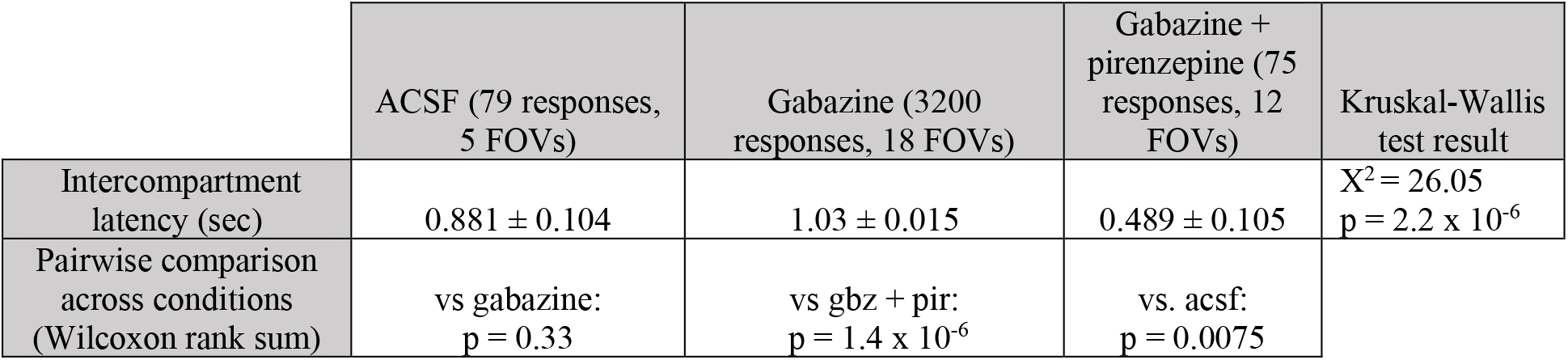
Intercompartment latency in ACSF, gabazine, and gabazine + pirenzepine Kruskal-Wallis test was used to test for overall drug effect on intercompartment latency between all conditions, with pairwise post hoc tests performed using Wilcoxon rank sum. Related to Fig 4D. Source data available in Figure 4-source data 1.

**Figure 4-figure supplement 6.**
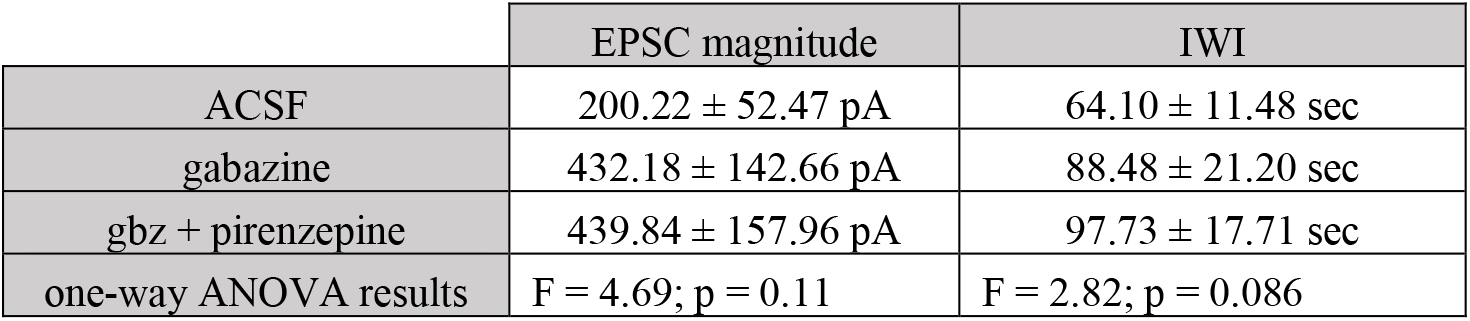
Wave properties in ACSF, gabazine, and gabazine + pirenzepine Wave properties are averages per RGC voltage clamp recording (n = 10 cells). Wave-associated EPSC and inter-wave interval (IWI) were compared using paired t-tests. Related to Fig. 4E. Source data available in Figure 4-source data 1.

**Figure 5-figure supplement 1.**
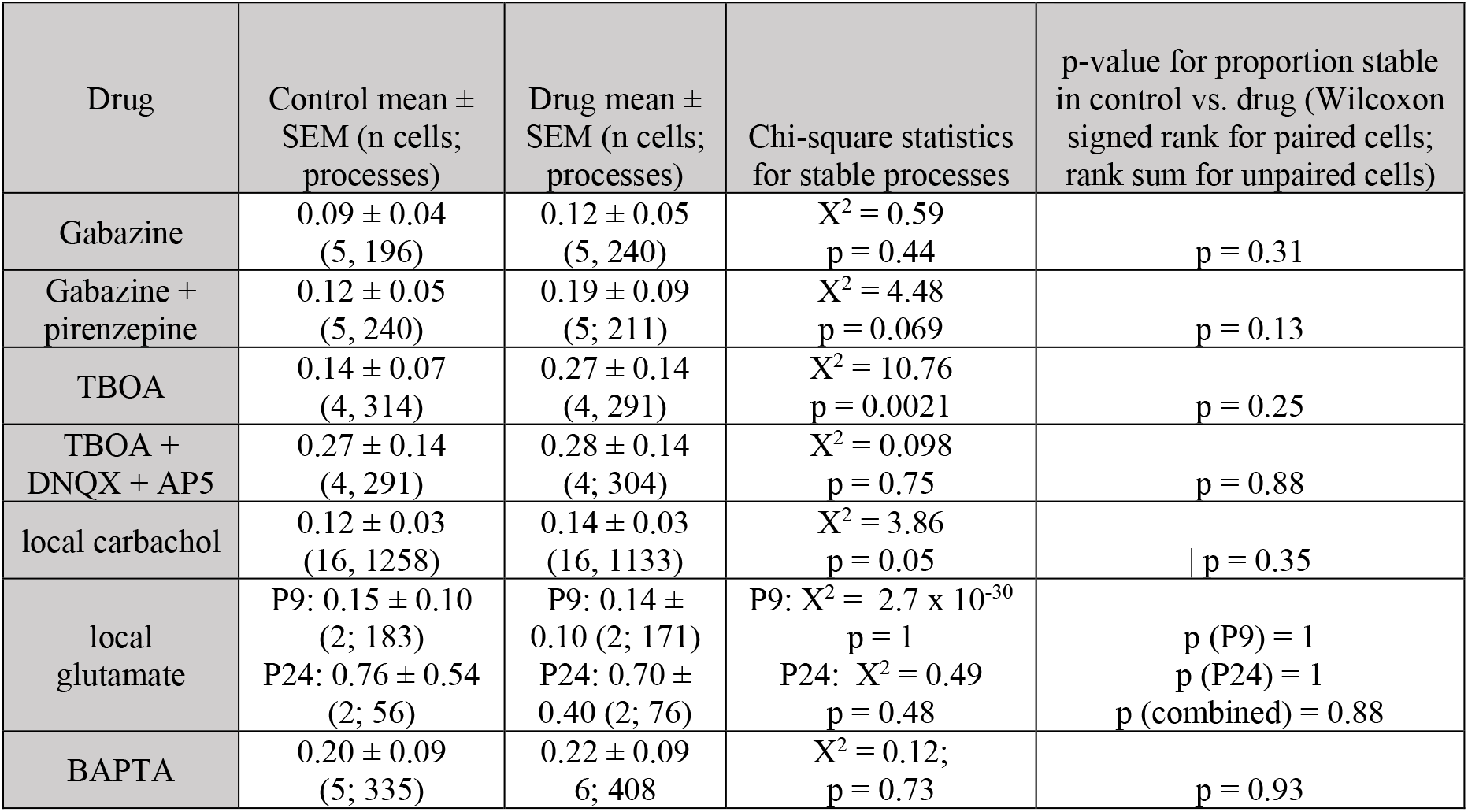
Comparisons of proportion stable processes across conditions Chi-square tests were used to test for differences in proportions of stable processes in control vs. drug using pooled count data. Pairwise comparison of proportion stable processes was performed using Wilcoxon signed rank test for paired cells and ranks sum test for unpaired cells. Related to Fig. 5F. Source data available in Figure 5-source data 1.

**Figure 6-figure supplement 1.**
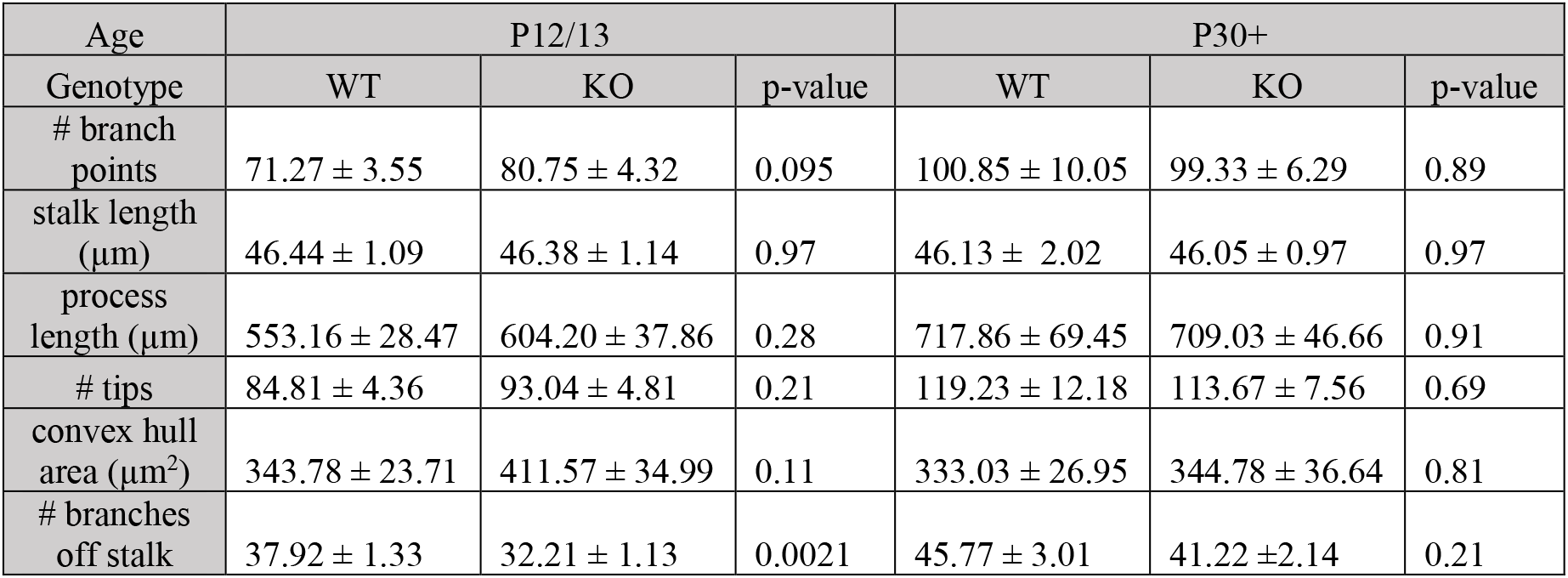
Comparisons of morphological properties between wild type and β2-nAChR-KO Müller glia Unpaired t-tests were used to compare morphological properties between wild type (WT) and β2-nAChR-KO retinas. Related to Fig. 6D. Source data available in Figure 6-source data 1.

